# Liver CYP4A autophagic-lysosomal degradation (ALD): A major role for the autophagic receptor SQSTM1/p62 through an uncommon target interaction site

**DOI:** 10.1101/2024.10.14.618315

**Authors:** Liang He, Doyoung Kwon, Michael J. Trnka, Yi Liu, Jade Yang, Kathy Li, Rheem A. Totah, Eric F. Johnson, A. L. Burlingame, Maria Almira Correia

## Abstract

The hepatic P450 hemoproteins CYPs 4A are typical N-terminally anchored Type I endoplasmic reticulum (ER)-proteins, that are inducible by hypolipidemic drugs and other “peroxisome proliferators”. They are engaged in the ω-/ω-1-oxidation of various fatty acids including arachidonic acid, prostaglandins and leukotrienes and in the biotransformation of some therapeutic drugs. Herein we report that of the mammalian liver CYPs 4A, human CYP4A11 and mouse Cyp4a12a are preferential targets of the ER-lysosome-associated degradation (ERLAD). Consequently, these proteins are stabilized both as 1%Triton X100-soluble and -insoluble species in mouse hepatocytes and HepG2-cells deficient in the autophagic initiation ATG5-gene. Although these proteins exhibit surface LC3-interacting regions (LIRs) that would target them directly to the autophagosome, they nevertheless interact intimately with the autophagic receptor SQSTM1/p62. Through structural deletion analyses and site-directed mutagenesis, we have identified the Cyp4A-interacting p62 subdomain to lie between residues 170 and 233, which include its Traf6-binding and LIM-binding subdomains. Mice carrying a liver-specific genetic deletion of p62 residues 69-251 (p62Mut) that includes the CYP4A-interacting subdomain also exhibit Cyp4a-protein stabilization both as Triton X100-soluble and -insoluble species. Consistently, p62Mut mouse liver microsomes exhibit enhanced ω- and ω-1-hydroxylation of arachidonic acid to its physiologically active metabolites 19- and 20-HETEs relative to the corresponding wild-type mouse liver microsomes. Collectively, our findings suggest that any disruption of CYP4A ERLAD results in functionally active P450 protein and consequent production of proinflammatory metabolites on one hand, and insoluble aggregates on the other, which may contribute to pathological aggregates i.e. Mallory-Denk bodies/inclusions, hallmarks of many liver diseases.

## Introduction

The physiological turnover of the integral mammalian endoplasmic reticulum (ER) proteins occurs via two major processes: ER-associated degradation (ERAD) involving ubiquitin (Ub)-dependent proteasomal degradation (UPD) (1–4) and/or ER-Lysosomal-associated degradation (ERLAD) entailing autophagic lysosomal degradation (ALD) (4–7), a process involving the concerted function of ≥ 30 autophagic (ATG) genes (8–13). In common with many integral ER-proteins, the physiological turnover of hepatic hemoproteins cytochromes P450 (P450s) also occurs via ERAD/UPD and/or ERLAD, henceforth denoted as UPD and ALD, respectively (14–26). Thus, while some P450s such as CYPs 3A incur largely UPD (14–17), others such as CYP2B1 incur largely ALD (18–20), yet others i.e. CYP2E1 incur both these processes (21–26), for reasons that remain yet to be elucidated. Nevertheless, while relative P450 preferences for each of these pathways normally exist, crosstalk between the two pathways exists with each pathway largely serving as a failsafe backup for the other, when the other is either overwhelmed or compromised, thus pre-emptying abnormal cellular protein accumulation and consequent proteotoxic stress (6,7, 27–32).

Initiation of the ALD process relies on a functional autophagy-related 5 (ATG5) protein, encoded by the *ATG5* gene (8–13, 33, 34). Upon activation by ATG7, ATG5 acts in concert with ATG12 and ATG16L1 to elongate the nascent phagophore or isolation membranes to give rise to the autophagic vesicles known as the “autophagosomes” (8–13). Our preliminary findings of proteomic analyses of phenobarbital (PB)-pretreated hepatocytes from ATG5-knockout (KO; ATG5^-/-^) mice and corresponding wild-type (WT; ATG5^+/+^) controls revealed that some P450s such as Cyps 1a2, 2a5 and 4a12a were relatively stabilized in ATG5^-/-^-mouse hepatocytes over those in corresponding WT-hepatocytes, revealing that most likely these P450s were preferential ALD-targets (*Supporting Information,* **Fig. S1**). By contrast, as expected, native P450s such as Cyp3a11 and Cyp2b10 that are known to preferentially incur UPD, were minimally stabilized in ATG5^-/-^ mouse hepatocytes. These proteomic findings also revealed that each stabilized P450 form consisted not only of “soluble” species solubilized by 1% Triton X-100, but also predominantly in the form of insoluble aggregates obtained as pellets upon 14,000g sedimentation of the 1% Triton X-100 extracts. Such aggregates required further solubilization by boiling with 8M urea/2M thiourea/4% CHAPS/Tris (TISO) buffer and upon immunoblotting (IB) analyses were shown to contain the parent P450 species as well as their high molecular mass (HMM) ubiquitinated and/or oligomerized species. Our proteomic finding that mouse liver Cyp4a12a is relatively stabilized upon ATG5-KO, suggested the plausible involvement of ATG5 in Cyp4a12a ALD (**Fig. S1**). Because very little is known about the P450 ALD process in general, we employed CYP4A proteins as model targets to mechanistically elucidate this process.

CYPs 4A are typical N-terminally anchored Type I ER-proteins, inducible by hypolipidemic agents i.e. clofibric acid and Wy-14,643, and other “peroxisome proliferators”, which are engaged in the ω-/ω-1-oxidation of various fatty acids including arachidonic acid, prostaglandins and leukotrienes as well as some therapeutic drugs (35–42). In this manuscript, we report that of the various human and mouse CYPs 4A, human CYP4A11 and mouse Cyp4a12a are preferential ALD targets. In addition to the ATG5 protein as well as other ALD-participant proteins, we also identify a major role for the autophagic receptor SQSTM-1/p62, a multifunctional protein scaffold, in their ALD (43–48). Furthermore, we also define an uncommon structural subdomain of p62 interacting with each of these two CYP4A proteins. This p62-subdomain resides within its N-terminal 170-233 residues and includes the Traf6-binding (TBS)-subdomain (228-233 residues) as well as at the least two other hotspots within its LIM-binding domain (LB). This p62 subdomain not only differs from its other well-known specific protein-interaction sites [Ub-association (UBA), microtubule-associated protein light chain 3 (LC3)-interacting region (LIR), Kelch-like ECH associated protein 1 (Keap1)- interacting region (KIR)] (43–59), but also excludes the positively charged R_183_/R_186_/K_187_/K_189_-patch located between its Zinc-finger ZZ- and TB-subdomains that we recently identified as a novel IκBα-interaction site (60).

## EXPERIMENTAL PROCEDURES

### Plasmids, vectors, and viruses

pcDNA6 and pcDNA3 vectors were from Invitrogen (Grand Island, NY). CYP4A11 and CYP4A22 were amplified by PCR with the templates of the human HepG2 cell cDNA, Cyp4a10, Cyp4a12a and Cyp4a14 were amplified by PCR with the templates of mouse primary hepatocyte cDNA. The CYP4A fragments were cloned into pcDNA3-Flag vector to generate pcDNA3-CYP4A10/Cyp4a12a/ Cyp4a14-Flag and pcDNA3-CYP4A11/CYP4A 22-Flag. Employing pcDNA3-CYP4A11-Flag and pcDNA3-Cyp4a12a-Flag as template, CYP4A11-Flag-T2A and Cyp4a12a-Flag-T2A fragments were obtained and cloned into ssfv-lenti-mCherry vector to generate ssfv-lenti-CYP4A11-Flag-T2A-mCherry and ssfv-lenti-Cyp4a12a- Flag-T2A-mCherry.

Plasmids pcDNA6-p62-myc, pcDNA6-p62-myc ΔZZ (128-163), pcDNA6-p62-myc ΔTB (225-251), pcDNA6-p62-myc N127, pcDNA6-p62-myc N224, pcDNA6-p62-myc N265, pcDNA6-p62-myc N320, pcDNA6-p62-myc N385, pcDNA6-p62-myc C225, pcDNA6-p62-myc C164, pcDNA6-p62-myc C104, pcDNA6-p62-myc K186A/K187A, and pcDNA6-p62-myc RRKK-A (R183R186K187K189AAAA) were constructed as previously reported (60). Truncation mutations were constructed by PCR cloning, or homologous recombination based NEBuilder® HiFi DNA Assembly kit (NEB, MA). Some p62 structural mutant expression plasmids were constructed specifically for the present study. The primers, templates, vectors used for the newly constructed plasmids are summarized **(Table S1,** *Supporting Information*):

Plasmids pAAV.TBG.PI.eGFP.WPRE.bGH and AAV.TBG.PI.Cre.rBG were gifts from Dr. James M. Wilson (Addgene viral prep # 105535-AAV8; http://n2t.net/addgene:105535; RRID:Addgene_105535 and Addgene viral prep # 107787-AAV8; http://n2t.net/addgene:107787 ; RRID:Addgene_107787). Lentiviruses ssfv-lenti-CYP4A11-Flag-T2A-mCherry or ssfv-lenti-Cyp4a12a-Flag-T2A-mCherry were packaged in house employing the protocol described below.

### Lentiviral packaging

HEK293T cells at a rapid replication state and grown to approximately 75%∼80% confluence in a 10 cm cell culture dish were passaged at 1:2 ratio for at least 2 consecutive days before seeding the cells for virus packaging. On day 0, HEK293T cells at 8.5∼9 x 10^6^ cells in a growth medium without antibiotics were seeded on a 10-cm culture dish to reach ≈ 90% confluence by the next day. They were then co-transfected with ssfv-lenti-CYP4A11-flag-T2A-mCherry or ssfv-lenti-Cyp4a12a-flag-T2A-mCherry along with pCMV-VSV-G and pCMV- DNF (ratio: 3:2:1, pCMV-DNF: 3 μg, pCMV-VSVG: 6 μg, ssfv-lenti: 9 μg) into HEK293T cells with the medium adjusted to 10 mL) with JetPEI transfection reagent (polyplus, jetOPTIMUS). Within 18 h, the old medium was (removed and replaced with fresh growth medium (10 mL). Twenty-four h after the transfections, mCherry-expression was monitored to determine >80-90% mCherry-expression efficiency. The medium was harvested 48 h later and replaced with fresh medium. Twenty-four h after harvesting the first batch of medium, the medium was harvested again. The pooled media containing the lentivirus were concentrated with Amicon Ultra-15 (UFC901024), aliquoted and stored at −80℃. Each frozen aliquot was thawed and used just once, avoiding repeated freeze-thaw-cycles that can lead to 50-90% loss of lentivirus.

### Liver-specific Atg5-KO mice

The ATG5-Floxed (ATG5 fl/fl) mice generated by Dr. N. Mizushima (Univ. of Tokyo) were provided by Dr. J. Debnath (UCSF). Liver-specific ATG5-KO were obtained through crossbreeding as follows: The breeding pairs were set as ATG5 fl/fl, +/+ and ATG5 fl/fl, Alb-Cre/+, such that 50% of the pups would be ATG5 liver conditional KOs (cKO) and the other 50% of pups, the WT controls. Liver tissues isolated from 6-week-old wild-type (WT, n = 3) and Atg5 KO (n = 3) mice were used for immunoprecipitation (IP) and/or IB analyses. The primers used for the tail-clip genotyping are indicated (**Table S2,** *Supporting Information*). Mice were fed a standard chow diet and maintained under 12-h light/dark cycle. All animal experimental protocols were approved by the UCSF/Institutional Animal Care and Use Committee.

### Liver-specific p62-mutant mice

Mice containing a liver-specific deletion of residues 69-251 of p62 (p62mutant, p62Mut) were generated as previously described (60). Alternatively, 4 week-old male p62 fl/fl mice were transduced with pAAV.TBG.PI.eGFP.WPRE.bGH (AAV8) (Addgene item #105535-AAV8) or AAV.TBG.PI.Cre.rBG (AAV8) (Addgene item #107787-AAV8) virus (1x 10^11^ vg each mouse) to generate WT (n = 4) and liver-specific p62Mut mice (a liver-specific deletion of residues 69-251 of p62, n = 5). Four weeks after the viral transduction, mice were killed and perfused livers used for microsomal preparations for IB analyses to verify the p62-deletion as well as functional and spectral assays. The primers used for the tail-clip genotyping are indicated (**Table S3,** *Supporting Information*).

### Primary hepatocyte culture

Hepatocytes were isolated from male C57BL/6 mice (8-week-old) by *in situ* liver perfusion with collagenase and purification by Percoll-gradient centrifugation by the UCSF Liver Center Cell Biology Core. Fresh primary hepatocytes (3.0 x 10^6^ cells/plate) were cultured on 60 mm Permanox plates (Thermo Fisher Scientific, Waltham, MA) coated with Type I collagen, and incubated in William’s E Medium containing 2 mM L-glutamine, Penicillin-Streptomycin, insulin-transferrin-selenium, 0.1% bovine serum albumin Fraction V, and 0.1 μM dexamethasone (DEX). After 6 h, Matrigel (Matrigel^®^ Matrix, Corning Inc. New York) was overlaid, and the cells were allowed to stabilize for 2 days.

### Quantitative RT-PCR Analyses

mRNA was extracted from the ATG5 WT/KO 8-week-old male/female mouse livers. TaqMan Universal PCR Master Mix (Applied Biosystems, Foster City, CA) was used to prepare a total of 20 µL of the PCR mix, with primers and probes at a final concentration of 909 and 125 nM, respectively. The amplification reactions were performed with an ABI Prism 7900 sequence detection system (Applied Biosystems, Foster City, CA) with initial hold steps (50 °C for 2 min, followed by 95°C for 10 min) and 40 cycles of a two-step PCR (92°C for 15 s, 60°C for 1 min). Each sample was measured in triplicate. The relative expression of each mRNA was calculated by the comparative method (ΔΔCt method, Ct = cycling time of the PCR amplifications). First, the mean Ct values of each Cyp4a and GAPDH for each sample were calculated from triplicate PCR reactions, and the ΔCt values for each sample were obtained by subtracting the mean Ct value of GAPDH from the mean Ct value of each Cyp4a. Then, the mean and standard deviation (SD) values for each group were calculated using the 2^−ΔΔCt^ values of samples in the same group. The qPCR primers used are listed (**Table S4**, *Supporting Information*):

### Degradation of Cyp4a in cultured hepatocytes

To induce Cyp4a, cultured mouse hepatocytes were pretreated with clofibric acid (clofibrate; 100 µM, Cat. No. 90323, Sigma-Aldrich Corp., St. Louis, MO) for 3 days. After the 3-day Cyp4a induction, cells were treated with the UPD inhibitor bortezomib (BTZ; 10 µM, Cat. No. 5.04314.0001, Sigma Aldrich Co.), or the ALD inhibitors 3-methyladenine (3-MA, 5 mM)/NH_4_Cl (50 mM) along with clofibric acid for 1 day. To induce autophagy, after the 3-day-Cyp4a induction, hepatocytes were treated with the mTOR inhibitor torin1 (1 µM, Cell Signaling, Cat. No. 14379) along with clofibric acid for 1 day. For cycloheximide (CHX)-chase analyses, cells pretreated with clofibric acid for 3 days, were treated with CHX (50 µg/mL) in the presence or absence of BTZ, 3MA/NH_4_Cl, or torin1 for up to 48 h. To monitor the Cyp4a degradation time course, cells were harvested at defined intervals and the lysates used for Cyp4a IB analyses.

### p62/NBR1-, ATG5- or Beclin1-KO HepG2 cells

The CRISPR/Cas9 system was used to knockout p62 or NBR1 or p62/NBR1 in HepG2 cell line. The CRISPR guide RNAs (gRNAs) were designed using CRISPR Design online tool (61). For p62, a sequence targeting exon 3 (TCAGGAGGCGCCCCGCAACA**TGG**) was used. For NBR1, a sequence targeting exon 1 (TTGGGCTGATATCGAAGCTA**TGG**) were used. Oligonucleotide containing the p62 or NBR1 CRISPR target sequence was annealed and ligated into Bbs1 linearized pSpCas9(BB)-2A-Puro (PX459) vector (Addgene #48139) to generate PX459-p62 and PX459-NBR1. HepG2 cells were transfected with PX459-p62 or PX459-NBR1 or co-transfected with both using X-tremeGENE HP transfection reagent (Roche) to generate single KO or double KO cells. The CRISPR/Cas9 system was also used to knockout ATG5 in HepG2 cell line. The CRISPR guide RNAs (gRNAs) lentivirus were purchased from Addgene. LentiCRISPRv2-ATG5 (Sequences targeting ATG5 exon 7 (CACCGGATGGACAGTTGCACACACT) was a gift from Dr. Edward Campbell (Addgene plasmid #99573; http://n2t.net/addgene:99573; RRID: Addgene_99573), and LentiCRISPRv2-Beclin1(Sequences targeting Beclin1 (CACCGATCTGCGAGAGACACCATCC) was also a gift from Dr. Edward Campbell (Addgene plasmid # 99574; http://n2t.net/addgene:99574; RRID: Addgene_99574). HepG2 cells were cotransfected using X-tremeGENE HP transfection reagent (Roche). Twenty-four h after transfection, cells were exposed to medium containing puromycin (5 μg/ml) and selected for 48 h, after which, cells were recovered with an antibiotic-free medium for 24 h before re-seeding into 100 mm dishes at 100 cells/dish. Cells were allowed to grow for 4-8 weeks until single cell colonies appeared. Single cell colonies were then picked and expanded for screening using Western IB analyses as guidelines.

### HepG2 and HEK293T cell culture and transfections

Cells were cultured as recently described (60). HepG2 cells, p62 KO HepG2 cells, p62/NBR1 KO HepG2 cells, NBR1 KO HepG2 cells, ATG5 KO HepG2 cells, and Beclin1 KO cells were cultured in minimal Eagle’s medium (MEM) containing 10% v/v fetal bovine serum (FBS) and supplemented with nonessential amino acids and 1 mM sodium pyruvate. HeLa, and HEK 293T cells (from ATCC) were cultured in DMEM (C11965500BT; Thermo-Fisher Scientific), supplemented with 10% FBS (10099-141C; Thermo-Fisher Scientific) and penicillin-streptomycin at 37°C with 5% CO2. For transfection experiments, cells were seeded on 6-well plates, when cells were 70% confluent, each cell well was transfected with 3 μg of plasmid DNA complexed with TurboFect transfection reagent (Thermo-Fisher, Grand Island, NY) for HEK293T cells, and X-tremeGENE HP transfection reagent (Roche, Indianapolis, IN) for HepG2 cells, according to the manufacturers’ instructions. At 40-72 h after transfection, cells were either treated as indicated or directly harvested for assays.

### Immunoprecipitation (IP) Analyses

Cell/tissue lysates (500 μg protein) or aggregates (200 μg protein) were incubated with 2% SDS at 75^○^C for 5 min, and then diluted 9-fold by volume with the cell lysis buffer. CYP4A antibody (5 μg, Abcam) was added, and incubated on a rocking platform at 4^○^C overnight. Pre-equilibrated protein G slurry (2 mg, Dynabeads^TM^ Protein G, Invitrogen) was then added, and the mixture incubated at room temperature for 3 h. The immunoprecipitated complexes were washed, eluted, and subjected to SDS-PAGE coupled with IB analyses against anti-Ub or p62 antibody. Normal rabbit IgG (Cat. No. 2729, Cell Signaling) was used as the negative control.

### Co-Immunoprecipitation (Co-IP) Analyses

Cells were transfected with indicated plasmids for 24-48 h and harvested with Cell Lysis Buffer (cat: #9803; Cell Signaling Technology, Inc. Danver, MA) or RIPA Buffer (cat: #9806; Cell Signaling Technology) supplemented with Halt™ Protease and Phosphatase Inhibitor Single-Use Cocktail (Cat: 78442). Cell lysates were sonicated for 10 s and then centrifuged at 13,000 rpm for 10 min at 4°C. Protein concentrations were determined by the bicinchoninic acid (BCA) assay. Cell lysates (1 mg) were then incubated with ChromoTek Myc-Trap® agarose beads (30 μL, Cat No: yta-20; Proteintech, IL) or ChromoTek GFP-Trap® agarose beads (30 μL, Cat No: gta-20; Proteintech, IL) at 4°C overnight, and then eluted by heating at 70°C for 10 min in 2X SDS-loading buffer. Eluates were subjected to IB analyses as described below.

### Immunoblotting (IB) analyses

Harvested hepatocytes and liver tissues were homogenized in cell lysis buffer (Cell Signaling Technology, Inc. Danver, MA) containg 10% glycerol, 5 mM N-ethylmaleimide, 1 μg/ml leupeptin, 1 mM, Na3VO4, 1 mM β-glycerophosphate, 2.5 mM sodium pyrophosphate and protease/phosphatase inhibitor (PI) cocktail. The homogenates were centrifuged (14,000xg, 10 min, 4 °C) and the supernatant was used as the cell lysate (Soluble fraction). The 14000xg pellet was solubilized in urea/CHAPS buffer (8 M urea, 2 M thiourea, 4% CHAPS, 20 mM Tris-base, and 30 mM DTT) and used as the “aggregate fraction”. Protein concentration was determined by the BCA assay and equal amounts of proteins [cell/tissue lysates (5 μg protein) or solubilized aggregates (5 μg protein)] were separated on 4-15% Tris-Glycine eXtended (TGX) polyacrylamide gels. Proteins were transferred onto 0.2 μ nitrocellulose membranes (BioRad, Hercules, CA) for IB analyses. Commercial primary antibodies were used for detecting CYP4A (rabbit monoclonal, Cat. No. ab140635, Abcam, Cambridge, UK), p62 (2C11, Abnova, Taipei City, Taiwan), LC3 (rabbit monoclonal, Cat. No. 12741, Cell Signaling Technology (CST)), Beclin-1 (CST, #3738), Atg5 (D5F5U) Rabbit mAb, CST, #12994), mCherry (E5D8F) (CST, #43590), DYKDDDDK Tag Antibody (Binds to the same epitope as Sigma’s Anti-FLAG® M2 Antibody) (CST, #2368) and ubiquitin (Ub; mouse monoclonal, Cat. No. ab7254, Abcam). β- actin (mouse monoclonal, Cat. No. A5316, Sigma-Aldrich) or GAPDH (D16H11) XP® Rabbit mAb (CST, #5174) was used as the protein loading control.

### CYP4A homology modeling

A CYP4A11 homology model was generated on the basis of the CYP4B1 crystal structure as previously detailed (62, 63) and has been available as supplemental File P450 – 4A11-homology-model.pdb (62). A mouse Cyp4a12a homology model was generated similarly to that of CYP4A11 (62), and a PDB file and the amino acid alignments of Cyp4a12a with those of the CYP4B1, CYP4A11 and other Cyp4a proteins are provided as *Supporting Information*.

### Mouse liver microsomal P450 spectral and Cyp4a functional assays

Liver from WT (N =4) and p62Mut (N = 5) mice were perfused with ice-cold 1.15% KCl, and liver microsomes were prepared and “washed” by rehomogenization/resedimentation in 1.15% KCl. Their spectral P450 content was determined in an aliquot (1.5 mg/mL) using a plate-based reduced carbon monoxide binding assay as described (64). Liver microsomal Cyp4a-dependent arachidonic acid ω-hydroxylase was monitored by the throughput UPLC-MS/MS quantitation method of total cytochrome P450 mediated arachidonic acid regio-isomers and cis/trans-EET metabolites (65), with the following exception: Because the two metabolites of interest, 19-HETE and 20-HETE, exhibit relatively short retention times of 2.31 and 2.55 min, respectively, and 4.11 min for the internal standard 12-HETE d8, a chromatographic separation of 9 min duration was employed with a mobile phase consisting of A (water) and B (acetonitrile) both containing 10 mM formic acid as follows: A mobile phase gradient of 0-6 min 65% A, 6-7.5 min 10% A, and 7.51-9 min 65% A. Results are expressed as Mean ± SD of N = 4 WT and 5 p62Mut ≈ 8 week-old male mice.

### In-Cell Chemical Crosslinking and Crosslinked Protein Capture for LC-MS/MS (XLMS)

For each crosslinking, thirty T150 flasks of cultured HEK293T cells were co-transfected with pcDNA3-CYP4A11[His]_6_ and pcDNA6-p62H181K/H190K/H192K-[HA]_3_ at a 4:1 w/w ratio for 48 h. The cells were treated with Bafilomycin (BAF) for 24 h before cell harvest. For chemical crosslinking, cells were washed with ice-cold PBS 3 times and collected in PBS (pH 9.0) with a cell scraper. DSS (disuccinimidyl suberate; 0.1 mM) was then added directly to the cell suspension, mixed gently by inversion and incubated on a rocker platform for 2 h in the dark at room temperature. Tris-buffer, pH 7.4, was then added to a final concentration of 20 mM to quench the crosslinking reaction. Cells were pelleted and then solubilized in 8 M urea (containing 0.02% cholate, 1 mM DTT, 20% glycerol, 1 mM AEBSF and other PIs) with sonication and cleared by centrifugation at 10,000 x g at 4°C to remove insoluble cell debris (pellet). The solubilized protein was mixed gently by end-to end rotation for 1 h at 4 °C with 8 mL of Hispur Ni^+2^-NTA-agarose (Thermo-Fisher, Cat: #88221) that had been equilibrated with 20 mM Tris (pH 7.9) containing 0.5 M NaCl, 10 mM imidazole, 0.02% cholate, and 20% glycerol to pull-down His-tagged CYP4A11[His]_6_ and crosslinked CYP4A11[His]_6_/p62[HA]_3_. The resin-protein slurry was poured into a column and washed 40 times with 8 mL 20 mM Tris (pH 7.9) containing 0.5 M NaCl, 20 mM imidazole, 0.02% cholate, and 20% glycerol (320 mL, total). His-tagged CYP4A11[His]_6_ and crosslinked CYP4A11[His]_6_/p62[HA]_3_ were eluted with 10 volumes of 20 mM Tris (pH 7.9) containing 0.5 M NaCl, 0.5 M imidazole, 0.25% cholate, and 20% glycerol (80 mL). The sample was concentrated on an Amicon Ultra 15 mL centrifugal filter (Millipore, UFC901096) to 1350 μl. Tris buffer (20 mM, pH 7.9) containing 0.5 M NaCl, 0.5 M imidazole, 0.25% cholate, and 20% glycerol, was added along with a 20% SDS containing loading buffer, heated at 70°C for 10 min. Eluates were then split into 50-60 gel lanes and subjected to SDS-PAGE and stained with Coomassie Brilliant Blue to visualize the bands for subsequent in-gel digestion. This purification resulted in an enrichment of crosslinked p62 and CYP4A11 species.

### IR Fluorescence Detection of DSS- or SIAB-crosslinked p62-CYP4A11 Species

Western IB analyses of the crosslinked p62-CYP4A11 complexes was first carried out with a two-color system using IRDye® 680RD goat anti-mouse IgG for labeling p62-[HA]_3_ and IRDye® 800CW goat anti-rabbit IgG for CYP4A11[His]_6_ which were then detected by IR fluorescence detection with an Odyssey® Fc Imaging System (LI-COR Biosciences).

### Mass Spectrometric (MS) Analyses

For in-gel digestion of *in cell* crosslinked samples, the indicated gel bands from 60 replicate lanes were processed as previously described (60) with the following modification: Digestion was performed at 37°C overnight using 400 ng of Trypsin/Lys-C mix (Promega) in a buffer of 50 mM ammonium bicarbonate with 0.01% ProteaseMax surfactant added (Promega). Following digestion, each replicate sample was individually desalted prior to pooling and drying on a speed-vacuum. Samples were resuspended in 0.1% formic acid for analysis on an Orbitrap Exploris 480 mass spectrometer (Thermo) coupled through an EASY-Spray nano ion source (Thermo) and a FAIMS Pro ion mobility interface (Thermo) to a Dionex UltiMate 3000 uPLC (Thermo) running an EASY-Spray column (75 µm x 50 cm column packed with 2 µm, 100 Å PepMap C18 resin; Thermo). The mobile phases were: Solvent A water/0.1% formic acid; solvent B: acetonitrile/0.1% formic acid. DSS-crosslinked sample was injected twice using different FAIMS compensation voltages (CV) with other parameters kept the same. The sample was loaded at 300 nL/min at 2% B, and then eluted with a 200 nL/min gradient from 2-25% B over 180 min. The column was then washed at 85% B and re-equilibrated back to 2%. The total run time was 237 min. Precursor ions were acquired from 375-1500 m/z in the Orbitrap (120k resolving power, 100% AGC, 50 ms max injection time). The FAIMS source was operated at 3 CVs (injection 1: −45, −60, −70 V, injection 2: −40, −50, −65 V) with product ions acquired for 1 sec cycle times at each CV. Precursor ions with charges 3-8+ and intensity greater than 50,000 were isolated in the quadrupole (1.6 m/z selection window) and dissociated by HCD with stepped 23, 30, 37% NCE. Product ions were acquired in the Orbitrap (30k resolving power, 100% AGC, 150 ms max injection time). 30s dynamic exclusion and the peptide monoisotopic ion precursor selection option were enabled.

Peaklists were generated using PAVA and proteins present in the sample were identified with Protein Prospector v6.5.2 with Trypsin specificity and 1 missed cleavage. Precursor and product ion tolerance were 10 and 20 ppm respectively. Carbamidomethylation of Cys was used as a constant modification. Variable modifications were: Met oxidation, loss and/or acetylation of protein N-terminal Met, peptide N-terminal Glu conversion to pyroglutamate, dead-end DSS modification at Lys and protein N-terminus, and incorrect monoisotopic peak assignment (neutral loss of 1Da). Up to 3 variable modifications per peptide were allowed. The SwissProt database (downloaded 2024_01_24) for Human, Bovine, Pig, *E Coli*, and Yeast was searched alongside a randomized database for FDR estimation. Additionally, the sequence of the p62 H181K/H190K/H192K triple mutant was manually included (63,290 target + 63,289 decoy sequences). Proteins were reported at a 1% FDR threshold and sorted by spectral abundance factor (SAF) (66).

### Statistical analysis

All results were expressed as Mean ± SD and analyzed by the two-tailed unpaired Student’s t-test. The acceptable level of significance was established at p < 0.05.

## RESULTS

### Mouse hepatic Cyps 4a normally turnover via both UPD and ALD

Cultured mouse hepatocytes were pretreated with the hypolipidemic drug clofibrate, a known Cyp4a inducer (35, 36, 67, 68), and then treated with bortezomib (BTZ), a diagnostic proteasomal inhibitor (69) which led to the stabilization of some hepatic Cyps 4a, consistent with their proteolytic turnover via UPD (**Fig. 1A**). Furthermore, BTZ, consistent with its proteasomal inhibition at a step beyond Cyp4a ubiquitination, also led to the dramatic accumulation of ubiquitinated Cyp4a species, as detected upon Cyp4a-immunoprecipitation (IP) and immunoblotting (IB) analyses with anti-Ub antibodies (**Fig. 1B**). Intriguingly, such Cyp4a IP also co-immunoprecipitated the autophagic receptor SQSTM1/p62 (**Fig. 1B**). Although, p62 is known to contain an Ub-association site (UBA) in its C- erminus (50), it is unclear whether native Cyp4a or its ubiquitinated species were the actual p62- targets. On the other hand, treatment with 3-MA, an inhibitor of the initial autophagic steps (10, 70), coupled with NH_4_Cl, a lysosomal inhibitor by virtue of its ability to alkalinize the intralysosomal pH, also led to the stabilization of the same or other hepatic Cyps 4a, consistent with their proteolytic turnover via ALD (**Fig. 1A**). That 3MA/NH_4_Cl indeed blocked ALD was evidenced by the accumulation of both the autophagosomal membrane marker LC3-II, the lipidated form of LC3-I (10–12), and of the autophagic receptor SQSTM1/p62, both documented ALD-substrates (10-12, 71; **Fig.1A**). That both UPD and ALD were involved in Cyp4a-proteolytic turnover was further evidenced by cycloheximide (CHX) chase analyses coupled with treatment with either diagnostic UPD or ALD inhibitor probes (**Fig. 1C**). These findings indicated that whereas BTZ stabilized Cyp4a-turnover at 24 h consistent with UPD targeting of mouse Cyp4a species/isoforms with the shorter lifespan, 3MA/NH_4_Cl stabilized Cyp4a-turnover at 48 h, consistent with ALD targeting of the longer lifespan Cyp4a species/isoforms (**Fig. 1C**). Collectively, these findings revealed that both clofibrate-inducible and/or constitutive CYPs 4a were turned over via both UPD and ALD pathways in cultured mouse hepatocytes (**Fig. 1**). Such ALD turnover of clofibrate-inducible Cyps 4a was enhanced upon cell treatment with Torin 1, an mTOR-inhibitor, known to stimulate protein ALD (72, 73; **Fig. 1D**). Consistent with Torin 1-mediated enhanced ALD, a parallel loss of both autophagic substrates p62 and LC3-II was also observed (**Fig. 1D**). CHX-chase analyses coupled with Torin-1 treatment revealed the marked acceleration of Cyp4a proteolytic turnover, consistent with Torin-1-elicited Cyp4a ALD stimulation (**Fig. 1E**).

**Fig 1.**
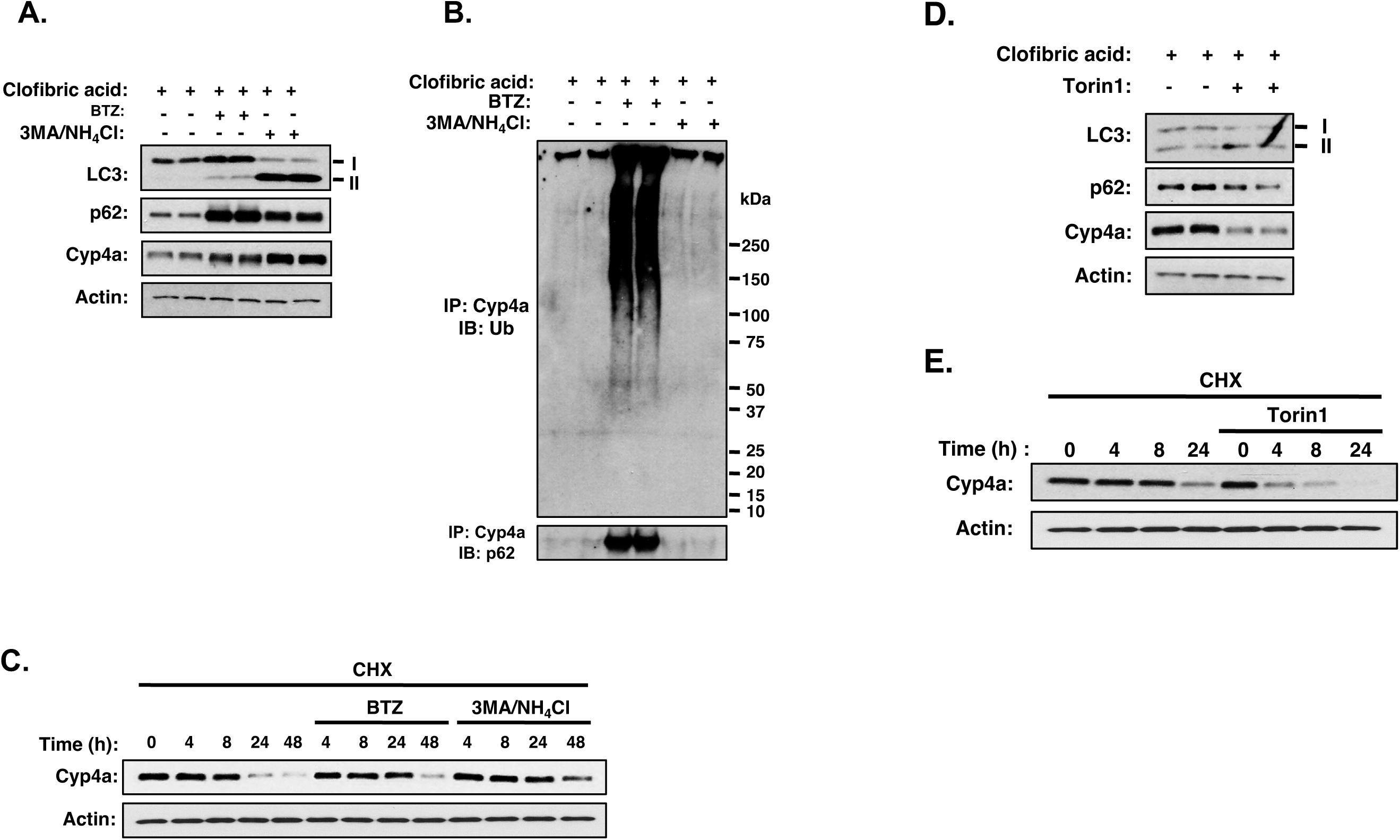
Cyp4a degradation via UPD and ALD in cultured mouse primary hepatocytes: Torin 1-elicited autophagic induction accelerates Cyp4a degradation. Cyps 4a were induced by a 3-day pre-treatment with clofibric acid. Mouse primary hepatocytes were isolated and cultured as detailed (Experimental Procedures). **A**. Cyp4a and autophagic flux as documented by the enhanced build-up of autophagic substrates LC3 and p62, upon hepatocyte treatment with dual 3MA/NH_4_Cl ALD inhibitors for 24 h. **B**. Cyp4a-IP of cell lysates upon treatment with UPD- and ALD-inhibitor probes reveals ubiquitinated Cyp4a species and p62-bound Cyp4a. **C**. CHX-chase analyses of clofibrate-inducible Cyp4a degradation. Cyp4a was induced by a 3-day pre-treatment with clofibric acid. BTZ (10 µM) and 3-MA (5 mM)/NH_4_Cl (50 mM) were employed as diagnostic UPD and ALD probes, respectively. Cyp4a levels were determined by IB analyses at 0, 4, 8, 24, and 48 h after treatment with CHX (50 μg/mL) in the presence or absence of BTZ or 3MA/NH_4_Cl. **D**. Torin 1 stimulates autophagic flux and Cyp4a ALD. Clofibric acid-pretreated cells for 3 d to induce Cyps 4a, were incubated for an additional day with clofibric acid with or without the autophagy activator torin1 (1 µM). The loss of p62 and LC3-II ALD-substrates was monitored as index of enhanced autophagy. **E**. CHX-chase analyses of clofibrate-inducible Cyp4a degradation accelerated upon Torin 1 treatment. Clofibric acid-pre-treated hepatocytes were incubated with CHX (50 μg/mL) with or without torin1 (1 µM) for 0, 4, 8, and 24 h, and Cyp4a content was determined by IB analyses.

To determine the physiological relevance of ALD to Cyp4a degradation, we employed liver conditional ATG5-KO mouse livers through crossbreeding ATG5 fl/fl with Alb-Cre mice (**Fig. 2A**), Consistent with the critical importance of ATG5-protein to liver protein ALD, upon ATG5-KO, autophagic impairment resulted in enhanced stabilization of hepatic LC3-I and LC3-II species, hepatic p62, as well as ubiquitinated hepatocellular proteins (**Fig. 2B**). Studies in these liver conditional ATG5-KO mouse livers, further substantiated our findings in cultured mouse hepatocytes (**Fig. 1**) by documenting hepatic Cyp4a stabilization upon genetic ATG5-KO (**Fig. 2C**). Notably, consistent with our earlier proteomic findings (**Fig. S1**), a nearly 2-fold stabilization of the 1%Triton-soluble Cyp4a species and an even higher, 3-fold stabilization of the 1%Triton-insoluble Cyp4a species (aggregates) were detected upon ATG5-KO of mouse livers relative to corresponding WT-controls (**Fig. 2C**). Cyp4a-IP of each P450-fraction coupled with Ub-IB analyses of the resulting immunoprecipitates revealed the presence of ubiquitinated and nonubiquitinated/native species (**Fig. 2D, E**). Intriguingly, Cyp4a-IP of lysates from ATG5-KO versus WT mouse livers coupled with p62-IB analyses of the immunoprecipitates, documented the relatively enhanced Cyp4a-p62 association upon ATG5-KO (**Fig. 2F**). This provided the very first clue that not only these two proteins plausibly interact intracellularly, but also that their interaction is detectably magnified upon ALD-impairment.

**Fig 2.**
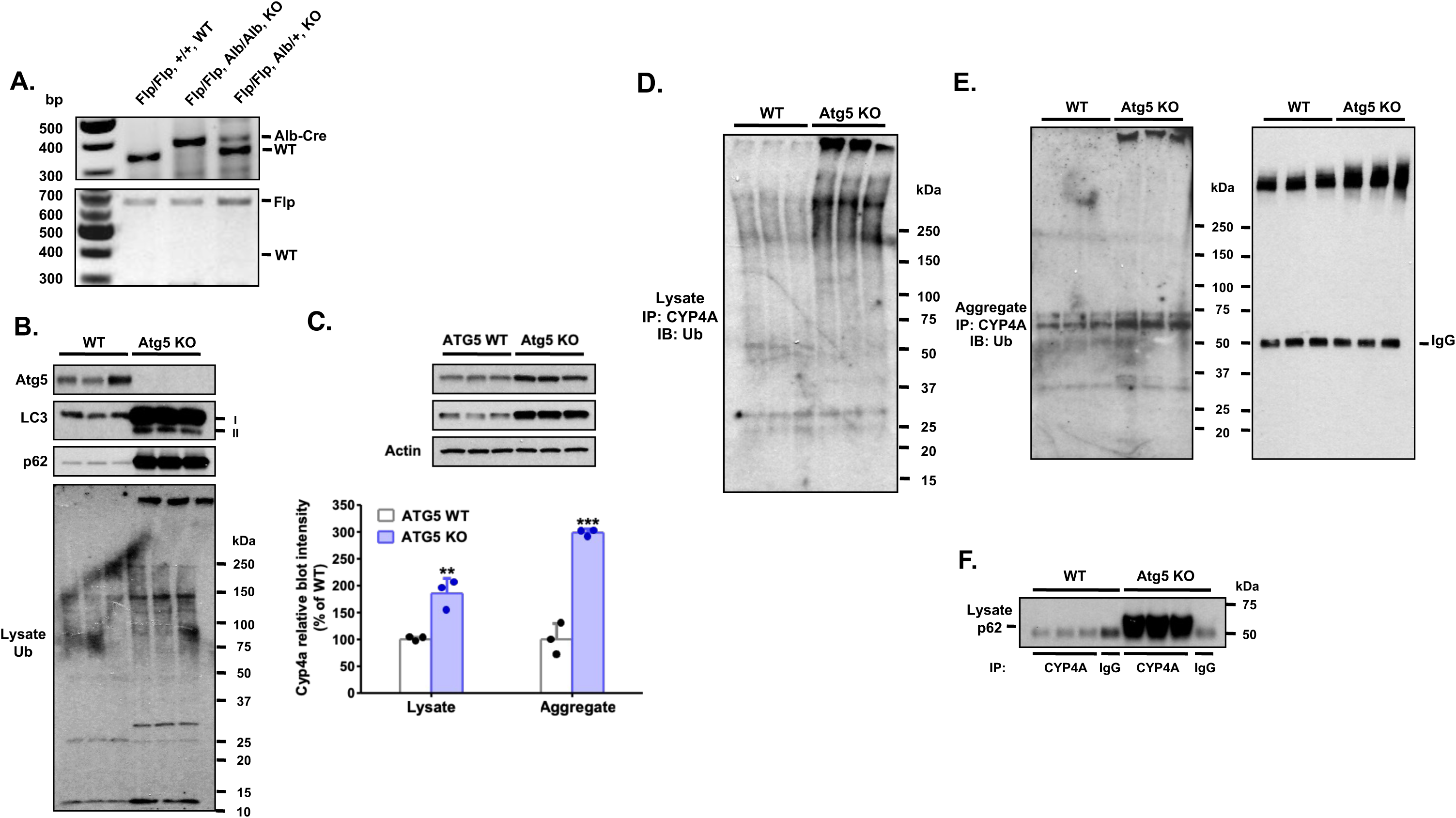
Relative accumulation of mouse hepatic Cyp4a levels of native and ubiquitinated protein species upon liver-specific Atg5-KO. **A**. PCR genotype of genomic DNA isolated via tail-clipping from WT and KO mice. **B**. Enhanced hepatic LC3-I and LC3-II as well as p62 levels and accumulated ubiquitinated proteins signal defective autophagic flux in Atg5 KO liver. **C.** Native Cyp4a levels were increased in soluble (lysates) and aggregate fractions of Atg5-KO mouse livers. Relative densitometric quantification (mean ± SD; N = 3 mice) of Cyp4a band in WT (100 ± 3.3%) vs ATG5-KO (185.6 ± 27.7%) lysates and of Cyp4a in WT (100 ± 29.9%) vs ATG5-KO (298.6 ± 7.1%) aggregates. **,***Significantly different from the corresponding WT mice (Student’s t-test, p < 0.01, and 0.001, respectively). Cyp4a in liver lysate or aggregate fraction was immunoprecipitated, and IB analyses were carried out using anti-Ub and p62 antibodies. Rabbit IgG was used as a negative control. **D**. IP analyses of Cyp4a in soluble liver tissue lysate followed by IB analyses with anti-Ub IgG reveals relatively enhanced accumulation of ubiquitinated Cyp4a. **E**. Cyp4a-IP of liver aggregate fraction reveals enhanced ubiquitinated Cyp4a in aggregate fraction of ATG5-KO mouse livers. **F**. Cyp4a-IP of soluble liver lysates coupled with IB analyses reveal enhanced association with co-immunoprecipitated p62 in ATG-KO mouse livers.

Mouse liver Cyps 4a comprise of at the least 3 different isoforms i.e. Cyp4a10, Cyp4a12a, and Cyp4a14, whereas CYP4A11 and CYP4A22 are the major human liver CYP4A isoforms (41, 74, 75). qRT-PCR analyses of mRNA extracted from male and female ATG5 WT mouse livers confirmed that Cyp4a12a is indeed the most abundant Cyp4a expressed in male mouse livers, followed in descending order by Cyp4a10 and Cyp4a14, whereas in female WT mouse livers the mRNA of Cyp4a14 and CYP4a10 predominate over that of Cyp4a12a (**Fig. 3**). Corresponding analyses of mRNA extracted from male and female ATG5 KO mouse livers, revealed that upon ATG5-KO, no significant changes in the expression of Cyp4a12a and Cyp4a14 mRNA, although some attenuation in Cyp4a10 mRNA expression was observed. These findings (**Fig. 3**) thus established that the enhanced Cyp4a content observed in ATG5-KO mouse livers was largely due to Cyp4a content stabilization upon ALD disruption rather than enhanced *de novo* protein synthesis.

**Fig 3.**
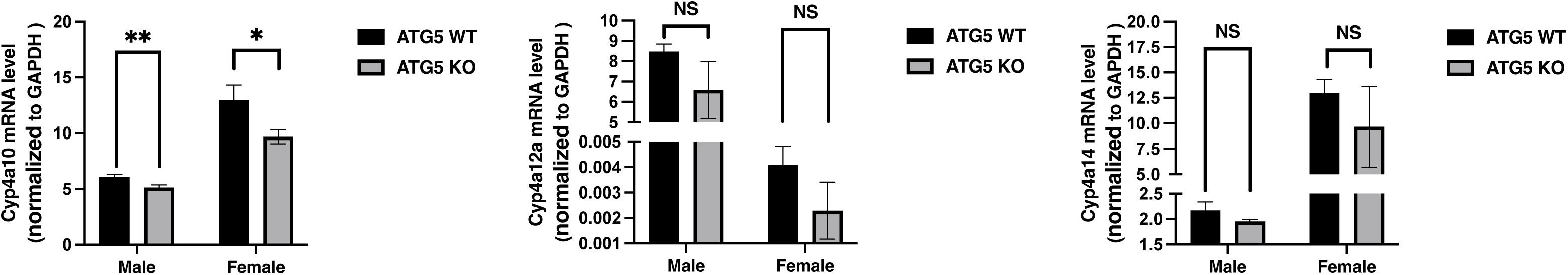
Cyp4a mRNA expression reveals that Cyp4a12a, the predominantly expressed Cyp4a in male mice, is unaffected upon ATG5 KO. qRT-PCR analyses were carried out with mRNA extracted from 8-week-old ATG5 WT/KO male or female mouse livers (N = 3 each) as detailed (Experimental Procedures). Each sample was assayed in triplicate. The relative gene expression of each P450 mRNA was calculated by the comparative method detailed in Experimental Procedures, after normalizing its threshold cycle (Ct) to that of GAPDH. Values are expressed as Mean ± SD. Statistically significant differences in these values were found between ATG5-KO and corresponding WT at p<0.01(**) and p<0.05 (*).

### Identification of human liver CYP4A11 and mouse liver Cyp4a12a as the principal ALD targets

Given our findings that mouse liver Cyp4a isoforms turn over via both UPD and ALD pathways, we sought to determine whether this duality was because our studies in mouse hepatocytes examined a mixture of hepatic Cyp4a isoforms, or whether each of these Cyp4a isoforms could be the target of both degradation pathways, much like CYP2E1 (21–26). To examine the proteolytic targeting of each individual Cyp4a isoform, we transduced HepG2 cells cultured in 12-well plates with lentivirus bearing a CYP4A/Cyp4a plasmid tagged with both FLAG and mCherry, with an intervening cleavable T2A peptide linker. Forty-eight h later, some cells were harvested as the 0 h controls, while the rest were treated with CHX, followed by BTZ, 3-MA/NH_4_Cl, Wortmannin (Wort) or DMSO as the vehicle control. Cells were cultured and some harvested at 20 h and others at 44 h. CHX-chase analyses of DMSO-treated cells revealed a major stabilization of CYP4A11 at 20 h and 44 h by 3-MA/NH_4_Cl and Wortmannin [a PI3K inhibitor that blocks autophagic initiation (10)], whereas the UPD inhibitor BTZ had only a minor effect (**Fig. 4A**). Similar findings were also obtained when lentivirus carrying a similar FLAG/mCherry-tagged Cyp4a12a was transduced into cultured HepG2 cells followed by a CHX-chase analyses and diagnostic UPD and ALD inhibitor probes, with the only difference being that BTZ stabilized Cyp4a12a to a greater extent than CYP4A11 at 20 h (**Fig. 4B**). These findings revealed that in contrast to CYP4A11, Cyp4a12a may be targeted to UPD to a slightly greater extent. CHX-chase analyses of both CYP4A11 and Cyp4a12a in cultured HepG2 cells revealed that consistent with their preferential ALD-targeting, Torin 1-treatment greatly accelerated their degradation (**Fig. 4C, D**). Parallel expression of Cyp4a10 and Cyp4a14 into HepG2 cells coupled with UPD and ALD diagnostic probes indicated that both proteins are largely UPD-targets (*Supplemental Information* **Fig. S2**).

**Fig 4.**
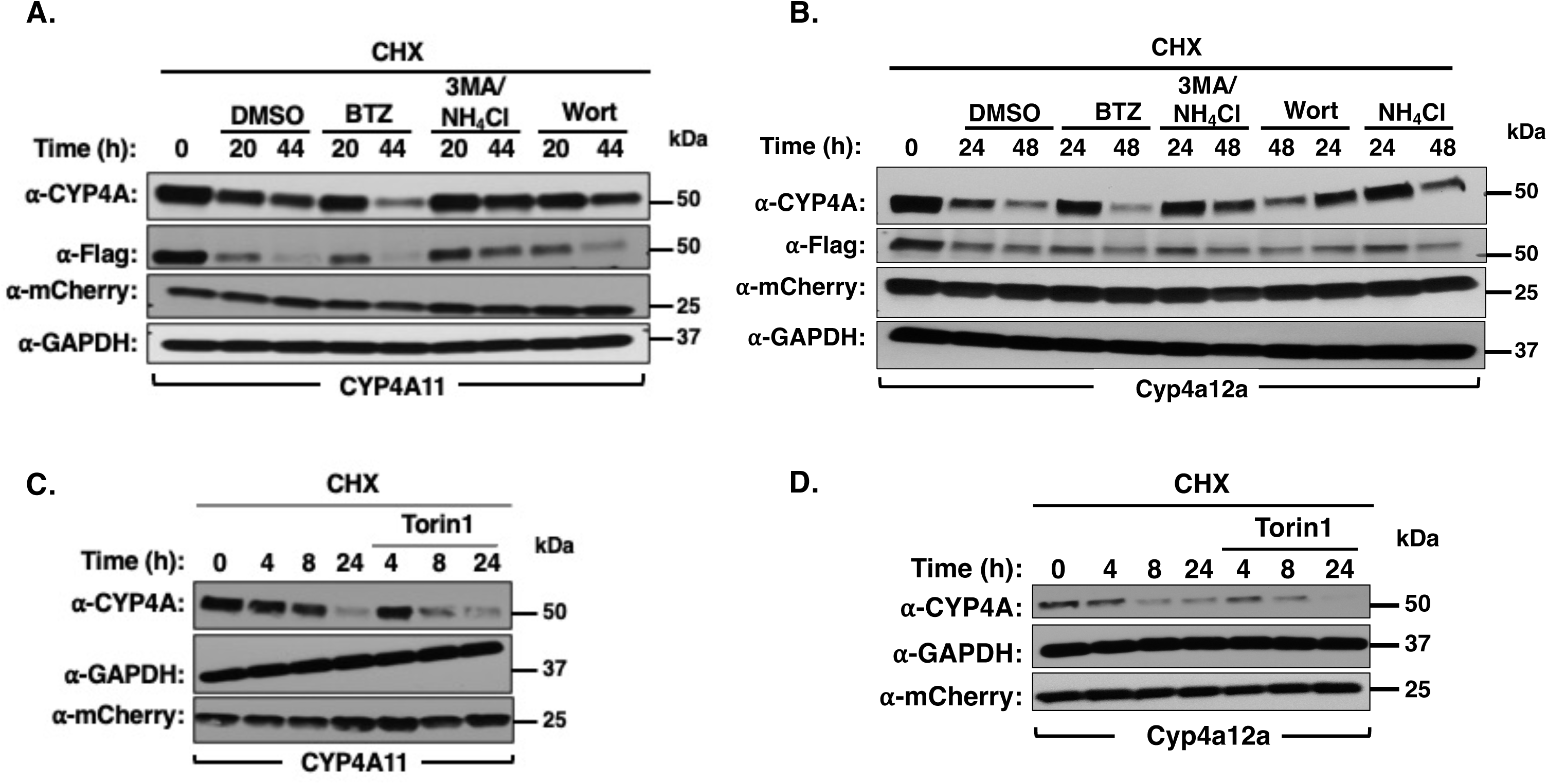
CYP4A11 and Cyp4a12a are preferentially degraded through ALD rather than UPD in HepG2 cells: Torin 1-elicited acceleration of CYP4A degradation. HepG2 cells were seeded in 12-well plates overnight, and then each well was transduced with either ssfv-lenti-CYP4A11-Flag-T2A-mCherry (**A**) or ssfv-lenti-Cyp4a12a-Flag-T2A-mCherry (**B**). Forty-eight h after transduction, cells were treated with CHX (50 μg/mL) for 10 min, and then the UPD-inhibitor BTZ, and ALD-inhibitors 3MA/NH_4_Cl or Wortmannin were added to specific wells, in parallel. Cells were harvested at indicated times (0, 20, and 44 h), and cell lysates (10 μg) subjected to IB analyses with CYP4A11 antibody or Flag antibody, with GAPDH employed as the loading control and mCherry as the transduction control. **C**, **D**. Cells transduced for 48 h as indicated in **A** and **B**, were subjected to CHX-chase analyses. At 0 h, Torin 1 (1 μM) was added to half of the wells and cells harvested at the indicated times, and lysates subjected to IB analyses with GAPDH employed as the loading control and mCherry as the transduction control. Representative immunoblots are shown.

Consistent with the above findings, lentiviral transduction of CYP4A11 or Cyp4a12a into WT and CRISPER-Cas-edited ATG5-KO HepG2 cells (**Fig. 5A**) showed relative CYP4A11 or Cyp4a12a protein stabilization in ATG5-KO cells relative to corresponding ATG5 WT (**Fig. 5B, C**). Furthermore, in HepG2 cells, similar KO of Beclin 1 (product of BECN1, the ortholog of yeast ATG6 and a key subunit of PI3K class III complexes involved in autophagic initiation; 9-13) (**Fig. 5A**), also resulted in CYP4A11 or Cyp4a12a protein stabilization upon viral transduction into BECN1-KO cells, relative to corresponding WT-controls (**Fig. 5D, E**). These findings revealed that the degradation of both CYP4A11 and Cyp4a12a rely on core autophagic initiation components.

**Fig 5.**
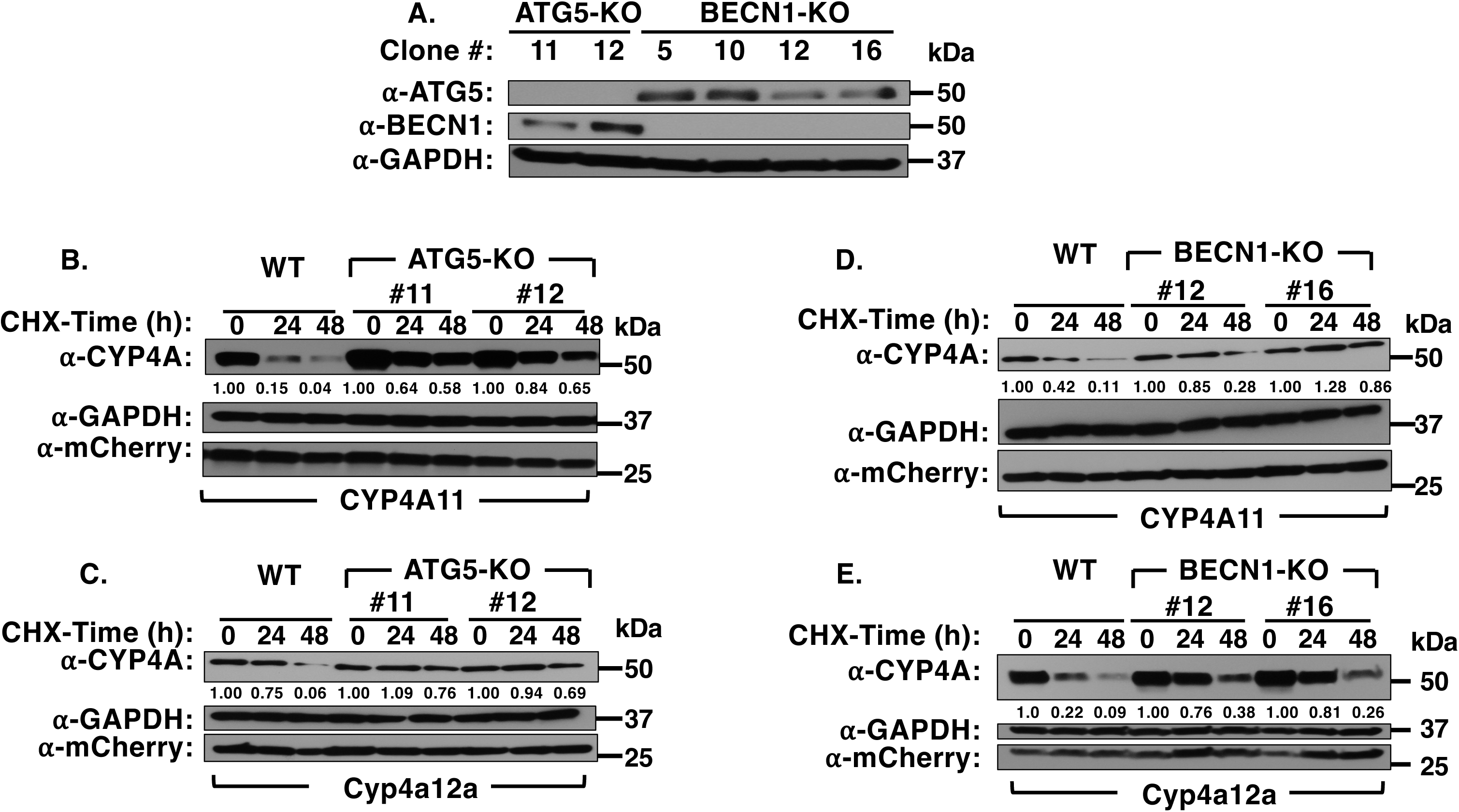
CYP4A11/Cyp4a12a protein stability upon ATG5/BECN1 KO in HepG2 cells: ATG5 KO and BECN1 KO HepG2 cells were generated through CRISPR/Cas9-mediated deletion of each gene (**A**). IB analyses of KO and corresponding WT cell lysates, probed with antibodies against ATG5 and BECN1 were carried out to assess the deletion extent of human ATG5 and BECN1 expression with GAPDH used as a loading control. WT, ATG5 KO HepG2 cells (**B**) and BECN1 KO HepG2 cells (**C**) were seeded in 12-well plates overnight, and then each well was transduced with either ssfv-lenti-CYP4A11-Flag-T2A-mCherry or ssfv-lenti-Cyp4a12a-Flag-T2A-mCherry. Forty-eight h after transduction, cells were treated with cycloheximide (CHX; 50 μg/mL). Cells were harvested at indicated times (0, 24, and 48 h). Cell lysates (10 μg) subjected to IB analyses with CYP4A11 antibody, along with GAPDH as the loading control and mCherry as a transduction control.

To further characterize the CYP4A ALD pathway participants in greater detail, we sought to enhance the detection of CYP4A-ALD interactants in cultured HEK293T-cells by a simultaneous Torin 1-elicited ALD-stimulation along with NH_4_Cl-mediated inhibition of lysosomal proteases (**Fig. 6A**). Cells were then co-transfected with a Flag-tagged CYP4A (4A11 or 4A22) or Cyp4a (4a10, 4a12a, 4a14) vector along with either an eGFP-N1 or eGFP-LC3-expressing vector for 48 h, followed by a 4 h-Torin 1/NH_4_Cl-co-treatment. Upon harvesting, cell-lysates were subjected to IP-analyses with a GFP-trap and subsequent IB analyses with anti-GFP IgGs and anti-CYP4A IgGs. By far, CYP4A11 was the protein co-immunoprecipitated with LC3 to the largest extent, followed by Cyp4a14 and Cyp4a12a. CYP4A22 was co-immunoprecipitated only to a very negligible extent with LC3, whereas Cyp4a10 was not at all (**Fig. 6A**). Given that LC3-II is a *bona fide* autophagosomal marker (10–12), these findings are entirely consistent with the finding that human CYP4A11 and mouse liver Cyp4a12a not only are largely targeted to ALD (**Fig. 4**), but also contain surface LC3-interacting region (LIR) motifs (**Fig. 6B**; 52), albeit not all readily accessible.

**Fig. 6.**
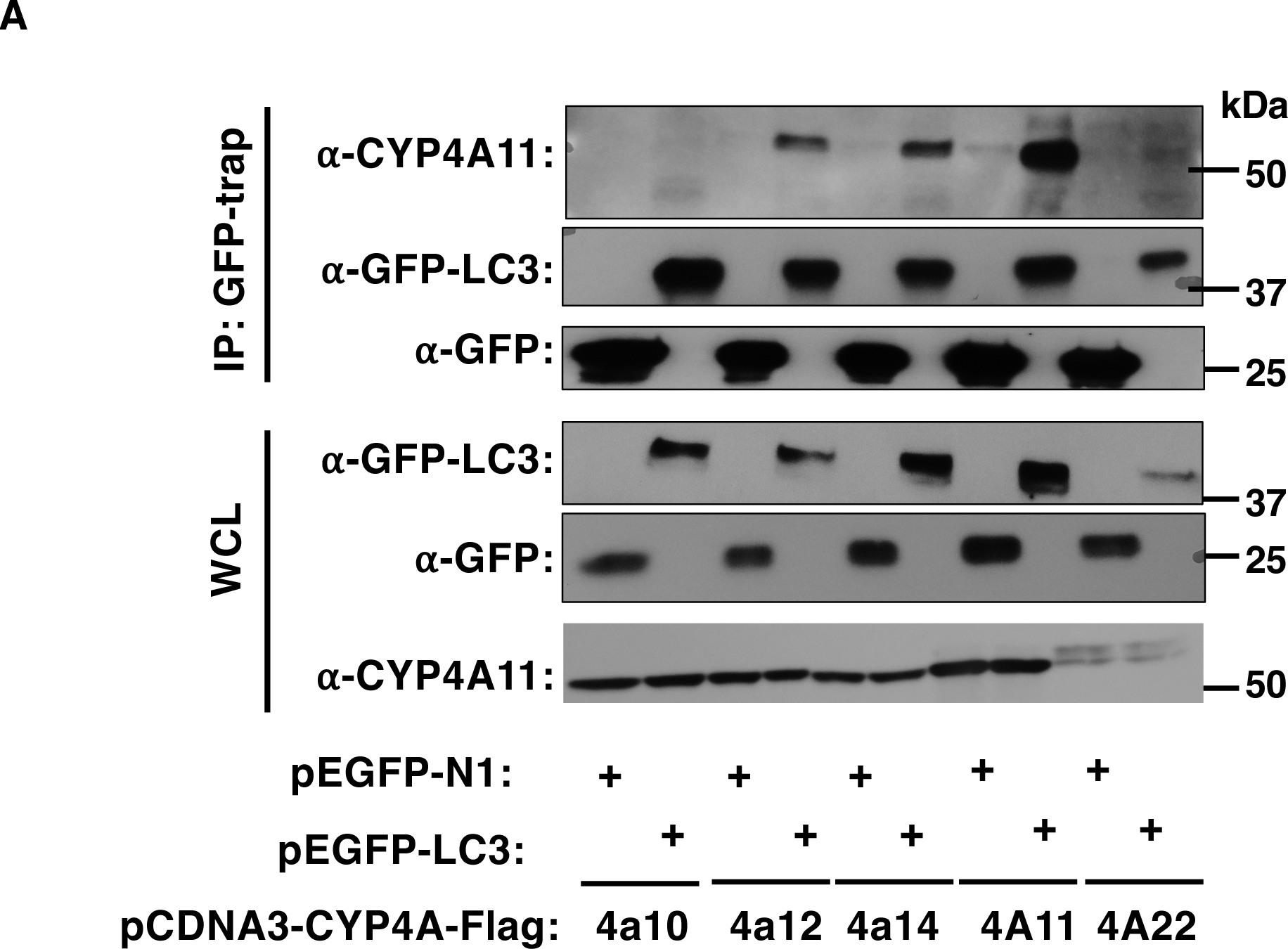
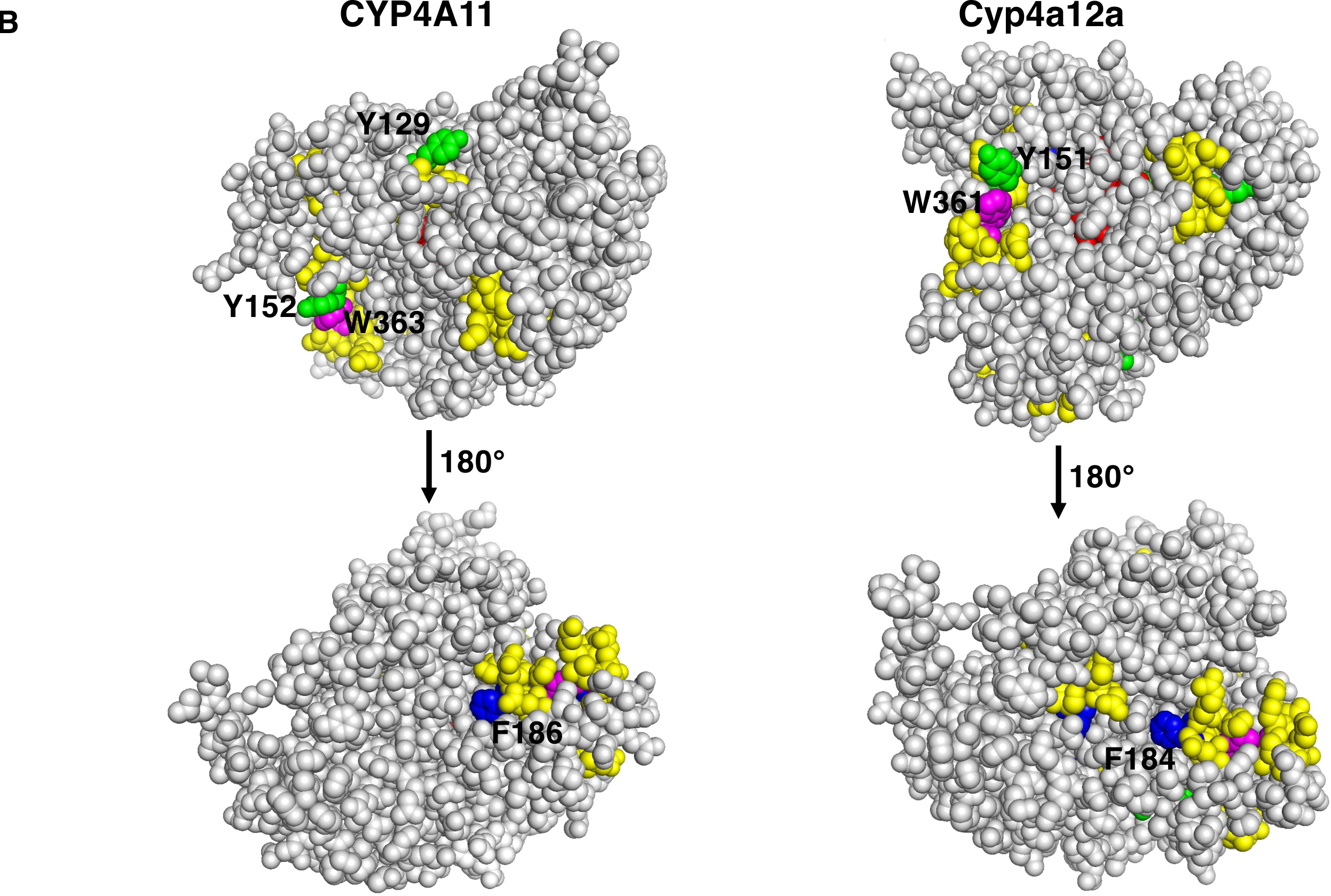
Relative LC3-interactions of CYPs 4A and depiction of CYP4A11 and Cyp4a12a external surface LIR motifs: **A.** HEK293T cells were seeded in 60 mm dishes overnight, and then each dish was co-transfected with GFP-LC3 (1.5 μg) and CYPs 4A (4 μg) for 48 h. Cells were treated with Torin (1 μM) together with NH_4_Cl (30 mM) 24 h prior to the cell harvest. The cells were harvested with cell lysis buffer containing Triton X-100 cocktail, and 0.1% SDS. Whole cell lysates (500 μg) were used for co-IP with Flag-beads (50 μL), followed by IB analyses with CYP4A11- and GFP-antibodies. IP, coimmunoprecipitation; IB, immunoblotting. **B.** Depiction of CYP4A11 and Cyp4a12a LIR motifs: PyMol depiction of P450 homology models were generated as previously described based on the structure of CYP4B1 (62, 63). LIR-motifs are relatively conserved 8-amino acid linear sequences (X_-3_X_-2_X_-1_W_0_X^1^X_2_LX_3_) required to dock at the hydrophobic Ub-like domain of the autophagosomal marker LC3 for autophagic targeting (52). In this motif, W_0_ has to absolutely be an aromatic (W, Y or F) residue, with a large hydrophobic (L, I, or V) residue at the L site, while X_-1_X_-2_X_-1_ and X_1_X_2_, may be, but not strictly, acidic D/E or phosphorylatable residues S/T. The full consensus LIR motif thus may consist of D/E/X-D/E/X-D/E/X-W/F/Y-D/E/X-D/E/X-L/I/V, and frequently reduced to the essential core sequence W/F/YXXL/I/V. Inspection of the CYP4A11-model reveals that although its primary structure contains at the least 10 LIR-motifs, only 4 (**GY_129_GLLLL, HY_151_DIL, EVF_186_QHVS and SITW_363_NHLD)** are on its surface, depicted in yellow with W (magenta), F (blue) and Y (green) residues. By contrast, although the Cyp4a12a primary structure contains 13 LIR motifs, only 3 (**HY_150_DIL, F_184_RHIT, SITW_361_NDLD)** are on its external surface and similarly color depicted. This may explain its relatively weaker interaction with LC3 compared to that of CYP4A11 (**Fig. 6A**).

### Plausible role(s) of cellular autophagic receptors in CYP4A11/Cyp4a12a ALD

Given our findings that these two P450s were not only the major targets of the cellular ALD pathway, but also interacted closely with the autophagosome marker LC3, we sought to determine whether this interaction was mediated by any of the common autophagic receptors p62, NBR-1 (neighbor of Braca 1 gene), TAX1BP1, NDP52, and OPTN (optineurin) (43, 57). In preliminary studies, WT HeLa cells and corresponding “penta-KO” cells upon knockout of these 5 genes (57) were transduced with either a CYP4A11- or Cyp4a12a-Flag-T2A-mCherry tagged viral vectors. After 48 h, a CHX-chase analysis of each P450 protein was carried out, with cells harvested at 0,16, 28 and 48 h (**Fig. S3**). These findings revealed that the penta-KO of these 5 autophagic receptors indeed impaired the degradation of both P450s. Because p62/NBR-1 are the major hepatic autophagic receptors, we then carried out a CRISPER-Cas-edited KO of these two major receptors, either individually or together in HepG2 cells (**Fig. 7**). CHX-chase analyses of each P450 protein revealed that both CYP4A11 and Cyp4a12a were stabilized in CRISPR-Cas9-edited p62/NBR-1 double-KO and p62-KO HepG2 cells, but not in NBR-1-KO HepG2 cells, thus indicating that p62 was most likely the major autophagic receptor involved in either CYP4A11 or Cyp4a12a ALD (**Fig. 7**). Co-transfection studies in HEK293T cells of each of the 5 P450 Flag-tagged pcDNA3 vectors with a p-Myc-tagged empty vector or p62-p-Myc-tagged vector, followed by p-Myc-pulldown and IB analyses, revealed that of the 5 P450s co-transfected, only CYP4A11 substantially interacted with p62, whereas Cyp4a12a and CYP4A22 did so albeit to a lower extent (**Fig. S4)**. A schematic representation of the relevant participants targeted in CYP4A11 ALD by the various diagnostic probes is shown (**Fig. 8**).

**Fig. 7.**
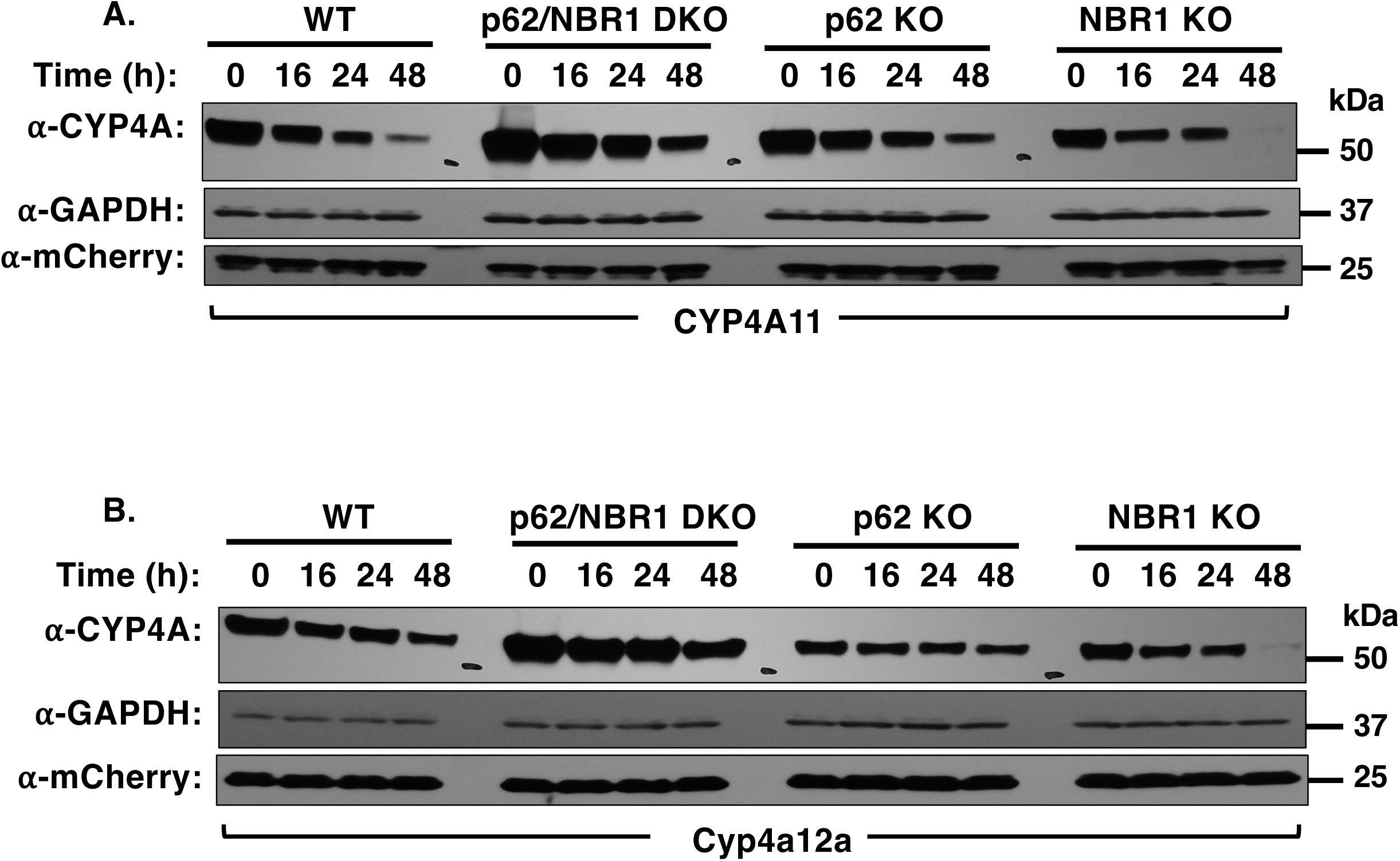
CYP4A proteins are stabilized upon p62 KO but not NBR1 KO of HepG2 cells: WT HepG2 cells, p62/NBR1 double KO, or p62 or NBR1 single KO HepG2 cells were seeded in 12-well plates overnight, and then each well was transduced with ssfv-lenti-CYP4A11-Flag-T2A-mCherry or ssfv-lenti-Cyp4a12a-Flag-T2A-mCherry. Forty-eight h after transduction, cells were treated with CHX (50 μg/mL). Cells were harvested at the indicated times (0, 16, 28, and 48 h). Cell lysates (10 μg) were subjected to IB analyses with CYP4A11 antibody, with GAPDH as the loading control and mCherry as the transduction control. CYP4A amounts relative to 0 h control were quantified from 3 experimental replicates and plotted. The half-life (t_1/2_) of CYP4A in WT and p62/NBR1 double KO, p62-KO and NBR-1-KO HepG2 cells was calculated based on a single exponential fit of the data with Graphpad Prism Version 6.07. They were found to be: CYP4A11 WT (21 h); p62/NBR1 double KO (39 h); p62-KO (31.8 h); NBR1-KO (24 h); Cyp4a12a WT (23 h); p62/NBR1 double KO (>48 h); p62-KO (>48 h); NBR1-KO (27.7 h).

**Fig. 8.**
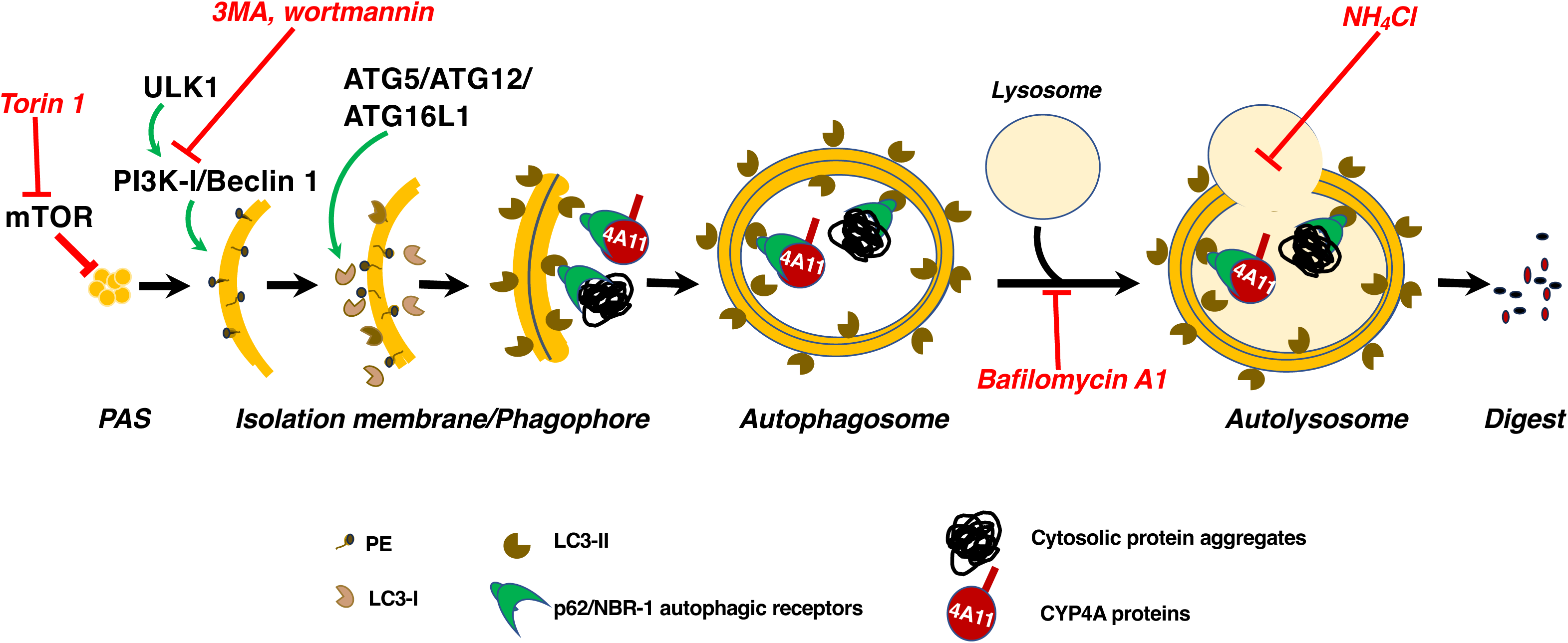
A schematic representation of CYP4A-ALD depicting the relevant autophagic participants targeted by the diagnostic probes employed and the sites of their inhibition in the autophagic process. Impairment at any stage in the autophagosomal biogenesis would result in CYP4A-aggregate accumulation. 3MA, 3-methyladenine; PAS, preautophagosomal structure; PE, phosphatidylethanolamine; PI3K, class III phosphatidylinositol 3-kinase (PI3K) complex I; ULK1, Unc-51-like Ser/Thr kinase 1.

### Identification of the CYP4A-p62-interaction site

The cellular autophagic receptor p62/SQSTM1 is a multifunctional protein scaffold known to interact closely with various cellular protein partners and/or motifs through well-defined structural elements and subdomains [Phox and Bem1p-domain (PB1), Zn-finger binding motifs (ZZ), LIM-domain interacting region (LB), TRAF6-binding (TB), LC3-interacting region (LIR), PEST1 and PEST2 motifs, Keap1-interacting region (KIR), and Ub-association region (UBA)] (43–56), and thereby to regulate various physiologically and pathophysiologically relevant processes. In addition to these previously well-established p62-interaction sites, we have recently identified a novel site for p62-interaction, stabilization and intranuclear transport of newly synthesized IκBα, that we found regulates the termination of NF-κB-transcriptional activation response (60).

Our findings of Myc-tagged p62-CYP4A11 co-transfection coupled with Myc-pulldown and IB analyses revealed not only the pull-down of CYPs 4A11 and 4a12a, but also upon film overexposure, of higher molecular mass (HMM) bands most likely reflecting either CYP4A oligomerization or CYP4A-ubiquitination, consistent with our earlier findings (**Fig. 2**). Thus, it was plausible that its UBA-domain could be a likely interaction site. To identify the p62-subdomains involved in its CYP4A-interaction, we carried out p62-deletion analyses through sequential deletion of various p62-plasmid regions corresponding to its known structural interacting elements and/or subdomains (43–56). Each of these Myc-tagged plasmids including the WT p62-Myc plasmid was co-transfected along with the CYP4A11- (or Cyp4a12a)-plasmid into HEK293T cells, followed by p62-Myc-IP and Western IB analyses of the co-immunoprecipitated CYPs 4A (**Fig. 9)**. These p62-CYP4A interaction analyses revealed that p62 constructs coding for residues 1-265, 1-320 and 1-385 were relatively more active than the parent p62, thereby excluding the p62 CT-domain [residues 266-440, which includes the Ub-interacting UBA-subdomain (389-434 residues)] as relevant to its CYP4A-interaction. Furthermore, the p62 construct coding for just p62 residues 1-127 exhibited a near loss of all CYP4A-interactions, whereas that coding for p62 1-224 residues was almost as defective. However, the PB-1 domain required for p62-oligomerization was found nonessential for the CYP4A-p62 interactions as all p62 constructs (coding for 164-440 and 104- 440 residues) devoid of the PB-1 subdomain, were fully competent in such interactions. Collectively, these p62 deletion analyses predicted its CYP4A-interaction domain to lie between residues 127-265. However, the deletion of just the ZZ-subdomain (residues 122-167) had little impact on the CYP4A-p62 interactions, but the deletion of the p62 TB-subdomain (residues 228- 233), reduced these interactions by ≈ 50%, thereby inferring the appreciable influence of this TB- subdomain on these interactions. Additional finer sequential deletion analyses identified the N-terminal p62 170-225 residues containing the intervening region (IR) between the ZZ- and TB- subdomains that is normally designated as the “LB-domain” as required for its cellular CYP4A11- interaction (**Fig. 9**). Further finer dissections of this p62 IR-subregion such as the deletion of residues 170-199 nearly abolished this interaction, while that of residues 170-179 or 190-199 greatly impaired the p62-CYP4A11 interactions. Surprisingly, neither the deletion of p62-residues 180-189 which contain the positively charged R_183_/R_186_/K_187_/K_189_ patch recently identified as a novel IκBα-interaction site (60), nor the mutation of these four residues to Ala failed to impair CYP4A11-p62 interactions. Collectively, the findings that individual deletion of the p62 TB-subdomain or that of residues 170-179 or 190-199 while failing to suppress these interactions greatly reduced them, suggested that most likely these p62-regions either represent tripartite hotspots for CYP4A11 interaction, or are structurally required for the optimal p62-recognition of CYP4A. Similar, but not identical findings were obtained upon similar deletion analyses of p62 and its Cyp4a12a interactions, suggesting that the p62-interaction hotspots for Cyp4a12a while residing in this general p62-subdomain, most likely involved different p62-residues and/or affinities.

**Fig. 9.**
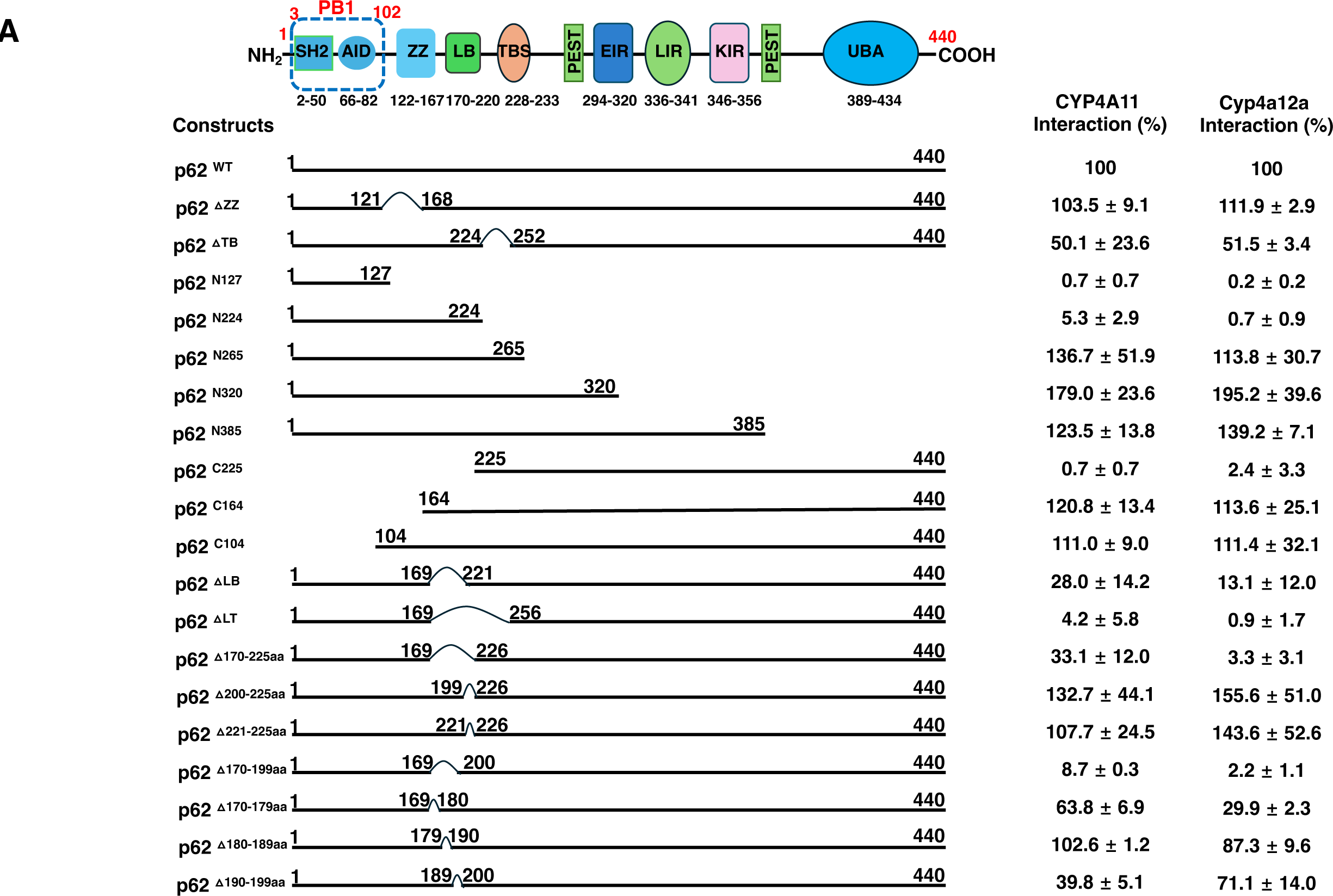
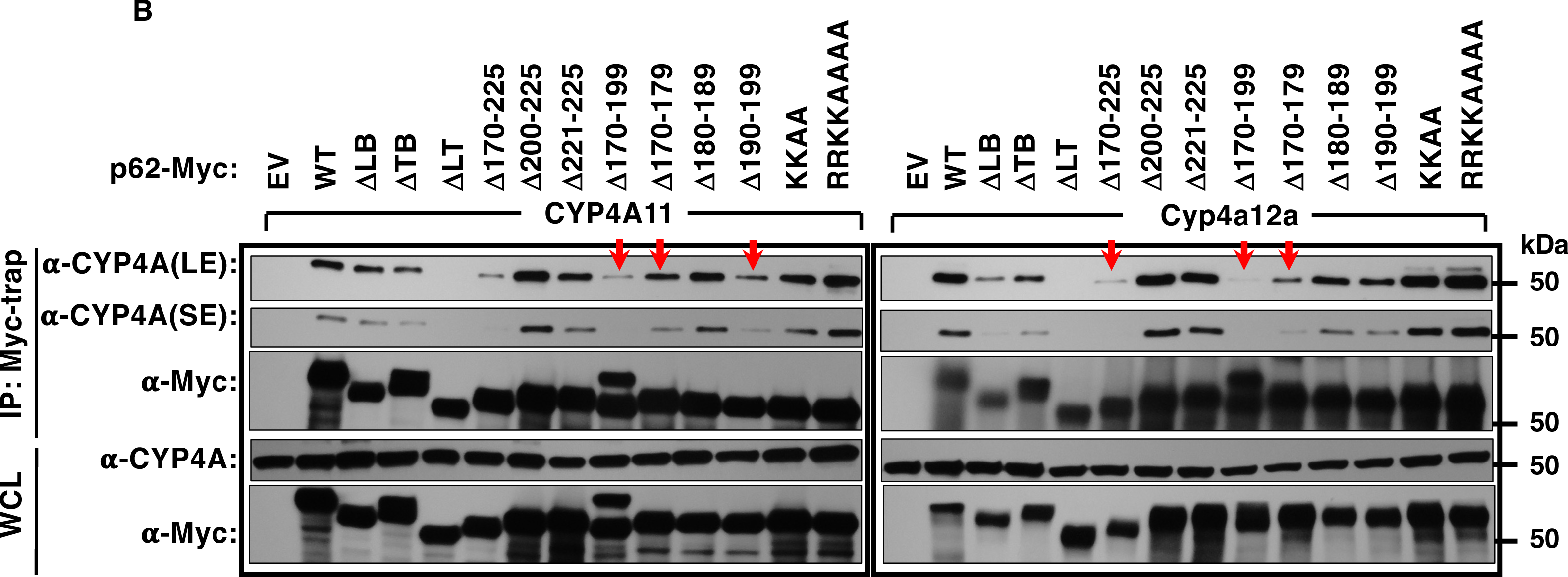
Identification of p62-CYPA11/Cyp4a12a-interaction domains through structural-deletion analyses of p62/SQSTM-1: Exclusion of positively charged R_183_R_186_K_187_K_189_. **A.** Various p62 deletion and truncation mutants were constructed as schematically shown. HEK293T cells were seeded in 60 mm dishes overnight, and then each dish was co-transfected with pcDNA3.1-CYP4A11 (4 μg) or Cyp4a12a (4 μg) and either WT or indicated mutant pcDNA6-p62-Myc vector (2 μg) for 48 h. Cells were treated with MG262 (1 nM)/NH_4_Cl (30 mM) 24 h prior to cell harvest. The cells were harvested with cell lysis buffer containing Triton X-100, protease and phosphatase inhibitor cocktail, and 0.1% SDS; Whole cell lysates (500 μg) were used for co- IP with Myc-trap (50 μL), followed by IB analyses with CYP4A11- and Myc-antibodies. Quantitative analyses of the CYP4A11 or Cyp4a12a interactions with various p62 deletion and truncation mutants relative to that with p62 WT expressed as 100%. Mean ± SD of at the least 3 individual experiments. **B.** A prototype immunoblot obtained in **A** is shown, additionally revealing that positively charged p62-residues R_183_R_186_K_187_K_189_ are not essential for CYP4A interaction. IP, coimmunoprecipitation; IB, immunoblotting.

To further identify the precise residues involved within this p62 170-199-residue region, we first deleted 20 residues 170-179 and 190-199 (Δ20aa) and found that the deletion of these two regions was almost as compromising to the p62-interactions with either CYP4A11 or Cyp4a12a as the deletion of the entire p62 170-199-residue span (**Fig. S5**). By contrast, mutation of the first 5 residues (170-174; Δ5A1) within p62 170-179 region resulted in very weak albeit detectable interactions relative to those of the parent p62 WT, whereas the deletion of just the next 5 residues (175-179; Δ5A2), albeit exhibiting weaker interactions than those with p62WT, were nevertheless appreciable. Similarly, deletion of the first 5 residues (190-194; Δ5A3) of the other relevant 190- 199-residue region, mitigated CYP4A-p62-interactions to a greater extent than the deletion of its next 5 residues (195-199; Δ5A4), although they were both weaker than those observed with p62WT. Collectively, these findings revealed that the plausible critical hotspots for CYP4A- interactions resided within these two regions, with some residues playing a more dominant role than others.

To further define the nature of these CYP4A-p62 interactions, identify the precise residues and gain some insight into their chemical nature, we generated Ala-mutations of single and/or multiple acidic/phosphomimetic (E), basic (H), phosphorylatable (S) residues within the 170-199 p62-subdomain through Ala-scanning mutagenesis (**Fig. 10**). Relative to the CYP4A11-interactions observed with p62WT, Ala-mutation of just E199 (M1) had no detectable effect (**Fig. 10A**), whereas Ala-mutation of various combinations of these residues mitigated CYP4A11-p62 interactions to various extents, with the greatest impairment observed with the combined Ala-mutation of 10 residues (M10: S170, H174, S176, S180, H181, S182, E177, H190, H192 and E199) (**Fig. 10B**). This impairment was just as severe as that observed with the deletion of the entire 30-residue (170-199) p62-subdomain. We found it instructive, that in contrast to Ala-mutants M6 and M7, both of which contained Ala-mutations of basic H190 and H192 residues and caused moderate impairments, M10 additionally contained Ala-mutations of S180, H181 and S182, which could also contribute to its severe impairment of the CYP4A11-p62 interactions. This combined ensemble of Ala-mutants also revealed that the basic H-residues were possibly more relevant to CYP4A11-p62 interactions than the acidic/phosphorylatable/phosphomimetic S/E-residues. Quite similar, but not identical Cyp4a12a-p62 interactions were also noted (**Fig. 10**).

**Fig. 10.**
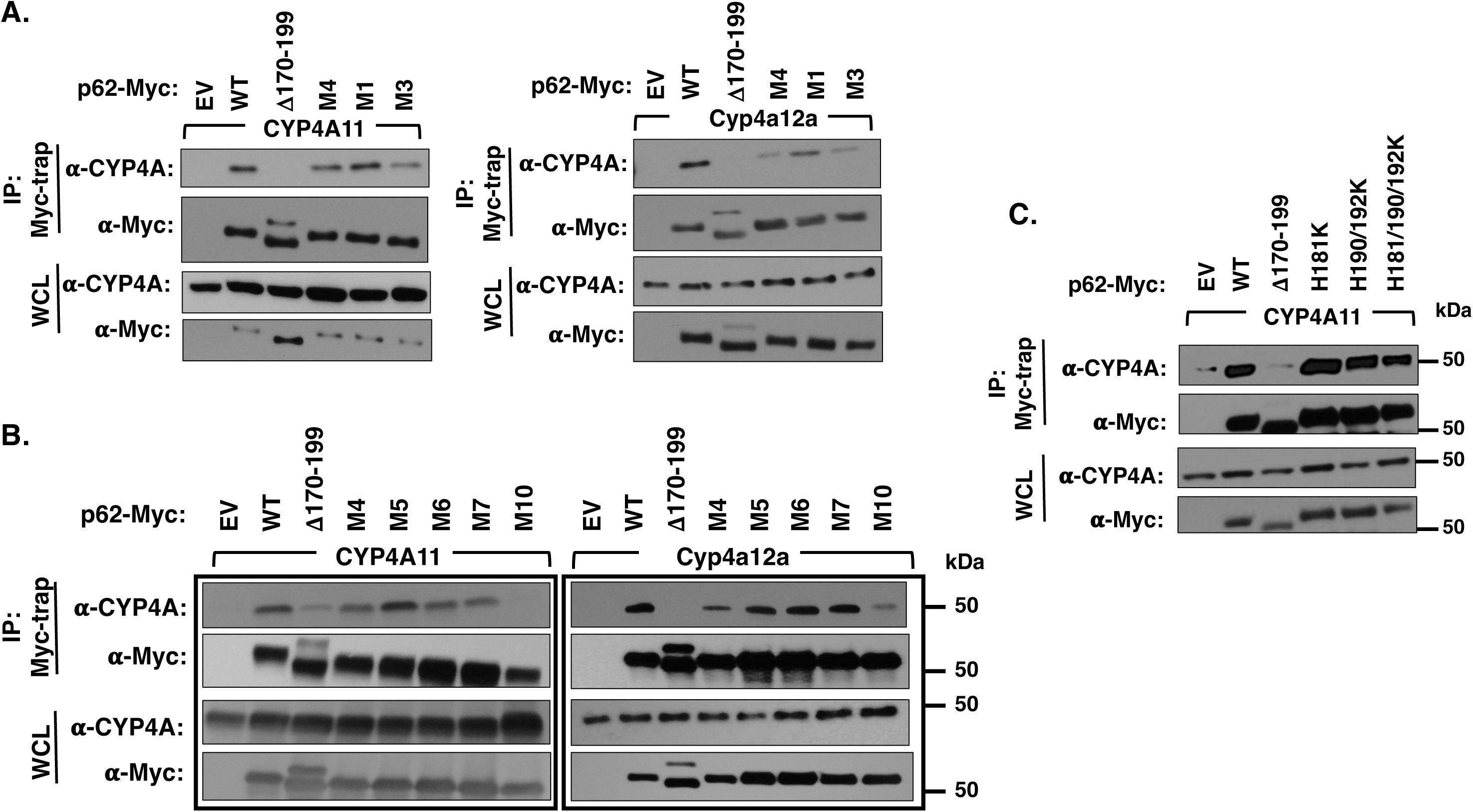
Identification of p62-CYPA11/Cyp4a12a-interaction domains through structural-deletion and site-directed mutagenesis analyses of p62/SQSTM-1. p62 subdomain-deleted or truncated, and site-directed mutants were constructed as schematically shown (**Fig. 9A**) or as indicated below. HEK293T cells were seeded in 60 mm dishes overnight, and then each dish was co-transfected with pcDNA3.1-CYP4A11 (4 μg) or Cyp4a12a (4 μg) and either WT or indicated mutant pcDNA6-p62-Myc vector (2 μg) for 48 h. Cells were treated with MG262(1nM)/NH_4_Cl (30 mM) 24 h prior to the cell harvest. The cells were harvested with cell lysis buffer containing Triton X-100, protease and phosphatase inhibitor cocktail, and 0.1% SDS. Whole cell lysates (WCL) (500 μg) were used for co-IP with Myc-trap (50 μl), followed by IB analyses with CYP4A11- and Myc-antibodies. IP, coimmunoprecipitation; IB, immunoblotting; EV, empty vector; WT, p62 WT vector; Δ170-199, p62 with deletion of 170-199 subdomain. **A.** CYP4A11 and Cyp4a12a interactions with p62 site-directed Ala-mutants M1 (E199A); M3 (S176A/E177A/E199A); M4 (S170A/S176A/E177A/E199A). **B**. CYP4A11 and Cyp4a12a interactions with p62 site-directed Ala-mutants M4 (S170A/S176A/E177A/E199A); M5 (S170A/H174A/S176A/E177A/E199A); M6 (S170A/S176A/E177A/H190A/H192A/ E199A); M7, (S170A/H174A/S176A/E177A/ H190A/H192A/E199A) and M10 (S170A/H174A/S176A/E177A/S180A/H181/ S182A/H190/H192A/E199A). **C**. CYP4A11 interactions with p62 site-directed Lys-mutants H181K, H190K/H192K, and H181K/ H190K/H192K.

To further characterize the plausible relevance of these basic H-residues in CYP4A11-p62 interactions, we examined specific p62-mutants containing K-mutations of residues H181, H190 and H192 (**Fig. 10C**). Indeed, p62 H181K-mutant greatly enhanced CYP4A11-p62 interactions, as did its H190K/H192K double-mutant and H181K/H190K/H192K triple-mutant over those observed with p62WT (**Fig. 10C**). We therefore exploited these super active p62 K-mutants in an attempt to define the precise hotspots of CYP4A-p62 interactions through DSS-chemical crosslinking/MS (XLMS) analyses, given that the cell-permeable chemical crosslinker DSS entails K-K intercrosslinks. Conceivably, this would enable us not only to identify the precise p62 hotspots involved, but possibly also the corresponding CYP4A11-interacting subdomains, given that unlike p62, P450s are not quite as amenable to structural deletion analyses.

### Identification of intracellular p62-CYP4A11 interactions through DSS-chemical crosslinking analyses

After initial optimization of the *in-cell* DSS-chemical crosslinking protocol (**Fig. S6**), we conducted DSS-chemical XLMS analyses in HEK293T-cells co-expressing p62[HA]_3_ and CYP4A11[His]_6_ (**Fig. 11A**). Upon *in-cell* DSS-crosslinking, the p62-CYP4A11[His]^6^ complex was isolated by passage of 8M urea-solubilized lysates onto a Ni^+2^-NTA-agarose column, extensive washing of non-specifically bound protein contaminants with 300-column volumes of 50 mM imidazole/1%cholate buffer, and elution with 500 mM imidazole/0.5% cholate buffers. This complex was however highly hydrophobic and tended to aggregate, requiring 8M urea for solubility and thus, intractable to any further enrichment through a secondary HA-IP. However, incontrovertible evidence that both CYP4A11 and p62 interacted intracellularly was obtained through the following: Upward mobility switch to >250 kDa of the crosslinked CYP4A11-p62 complex from their individual masses (i) upon SDS-PAGE *(denoted by black arrow* ***Fig. 11D***); (ii) two-color Western IB analyses of the DSS-crosslinked p62 and CYP4A11-species followed by LI-COR Odyssey IR fluorescence detection (**Fig. 11B & C**) as visualized by the yellow merged band (*marked by an yellow arrow,* ***Fig. 11C***) upon overlap of the immunoblot fluorescence signals detected with either IRDye 680RD goat anti-mouse IgG (p62[HA]_3_) or IRDye 800CW goat anti-rabbit IgG (CYP4A11[His]_6_); and (iii) LC-MS/MS analyses of the Lys-C/Trypsin in-gel digests of the SDS-PAGE band *(denoted by black arrow* ***Fig. 11D***) which verified the presence of p62 (38 unique peptides and 84.8% peptide coverage) and CYP4A11 (60 unique peptides and 82.1% peptide coverage) (*Supporting Information, Table S5*). However, despite our exhaustive attempts to optimize the recovery of the digested DSS-crosslinked peptides by inclusion of the Protease MAX surfactant (Promega) during the in-gel digestions, the high aggregation proclivity of the highly hydrophobic p62-CYP4A11 complex prevented detection of any DSS-crosslinked individual peptides. The parallel detection of DSS-crosslinked peptides of other irrelevant intracellular proteins, nevertheless provided evidence that DSS functioned *in vivo* as it should (*Supporting Information, Table S5*). Thus, although we could definitively attest to intracellular CYP4A11-p62 interactions and identify the p62-interaction subdomains through structural deletion/site-directed mutagenesis analyses, the CYP4A11 interacting hotspots remain yet to be defined.

**Fig. 11.**
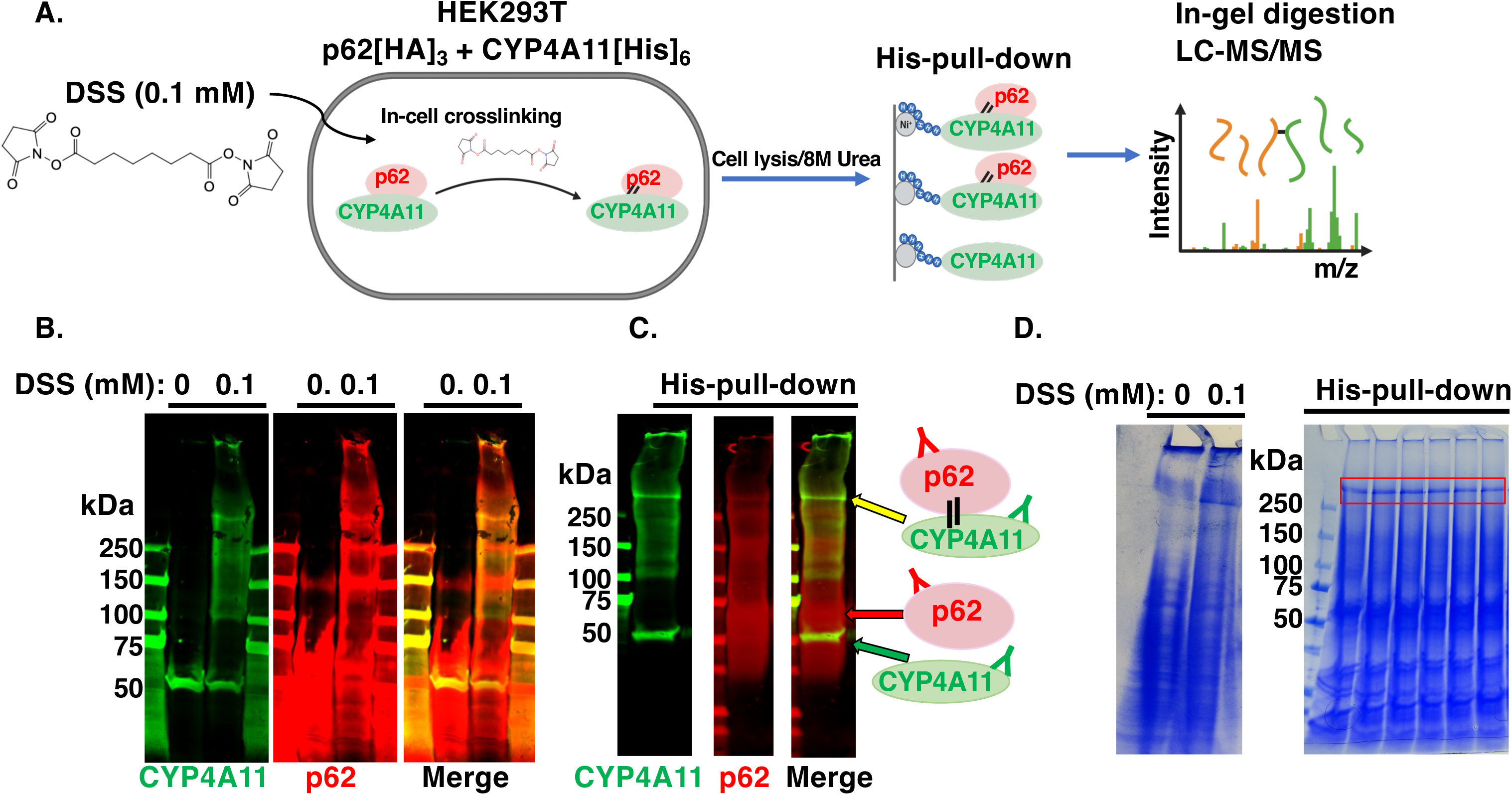
p62–CYP4A11 interactions identified by *in-cell* chemical crosslinking and mass spectrometric analyses. **A**. Schematic workflow for p62-CYP4A11 *in-cell* DSS-chemical XLMS analyses. **B**. Representative Western IB analyses of p62-[HA]_3_ and CYP4A11[His]_6_ before and after *in-cell* chemical crosslinking with 0.1 mM DSS and HisPur Ni^+2^-NTA protein purification. Crosslinked species were indicated by yellow arrow. **C**. Representative SDS-PAGE from large scale *in-cell* chemical crosslinking with 0.1 mM DSS and HisPur Ni^+2^-NTA protein purification. Bands excised for in-gel digestion and LC-MS/MS are demarcated by the *red box and indicated by the black arrow*.

### Physiological relevance of p62-Cyp4a12a interaction domain to Cyp4a12a ALD

To further verify the physiological relevance of intact p62-Cyp4a-interactions *in vivo*, we employed a mutant mouse carrying a liver-specific genetic ablation of the p62 residues 69-251 (p62Mut; 60). This p62Mut thus lacks the critical hepatic CYP4A-p62-interacting subregions (LB-residues 170-190, and TB-subdomain residues 228-233) (**Figs. 9 & 12A**).

**Fig. 12.**
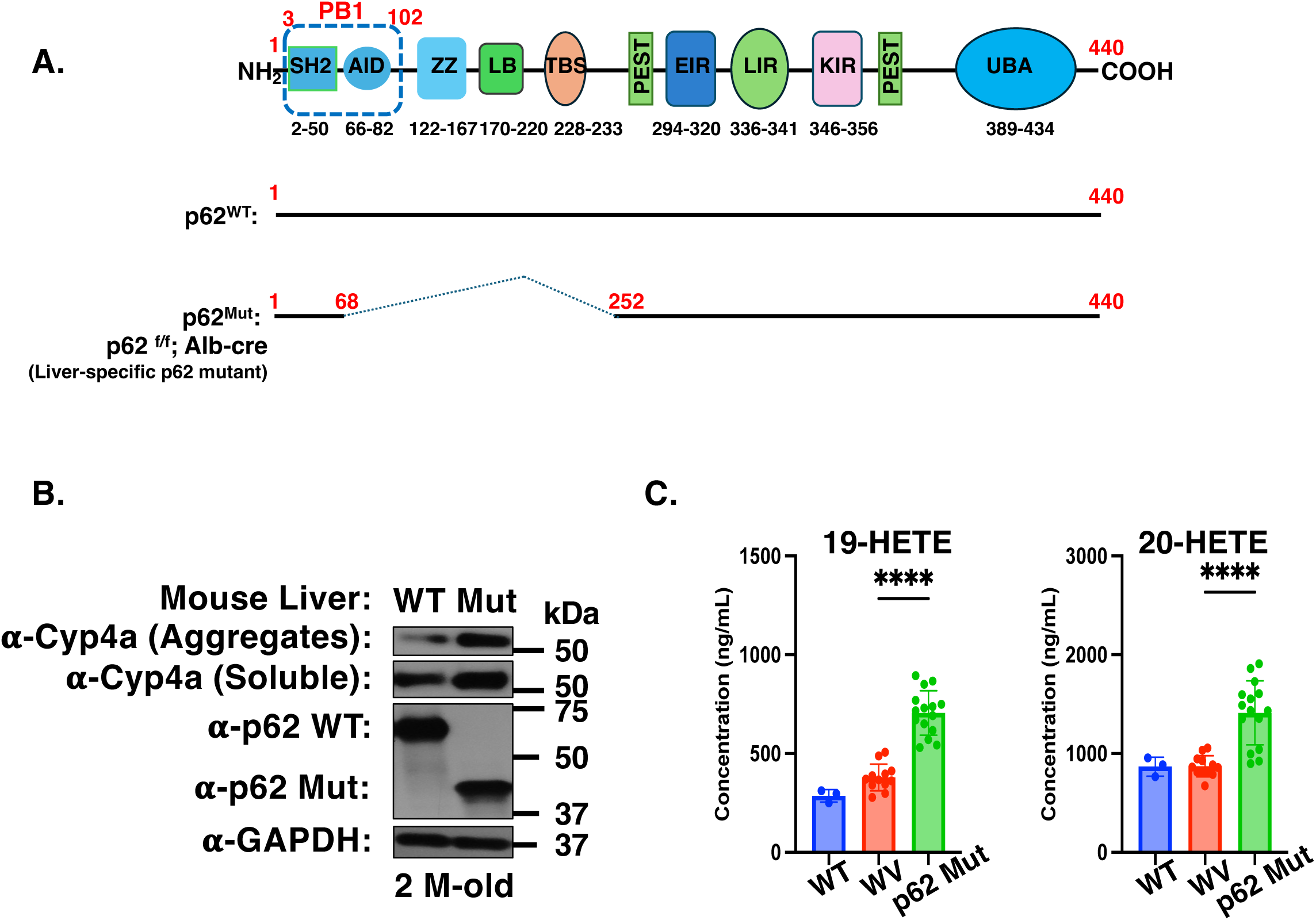
Cyp4a levels in p62Mut vs WT mouse livers and cultured hepatocytes: Functional Cyp4a stabilization in p62Mut-livers. **A.** Schematic depiction of the p62-region deleted in the liver-specific p62Mut mice. **B.** p62 Western IB analyses of liver lysates isolated from 2-month-old p62 WT and p62Mut mice. Hepatocytes were homogenized in cell lysis buffer and sedimented at 14,000g to obtain the insoluble CYP4A aggregates. The pellet containing CYP4A aggregates was solubilized in TISO buffer as described (Experimental Procedures). Cell lysates or solubilized pellet (10 μg) were subjected to IB analyses with CYP4A11 antibody, using GAPDH as the loading control. **C.** Liver microsomes isolated from 2-month-old p62 WT and p62Mut mice, were subjected to P450 spectral determination by the reduced-CO-reduced spectral assay, and to functional assay of arachidonic acid ω-hydroxylated metabolites 19- and 20-HETEs, as described (Experimental Procedures). Liver microsomal P450 content (Mean ± SD of untreated 2-month-old p62 WT and p62Mut mice was found to be 1.1 ± 0.064 (N = 4), and 1.4 ± 0.22 (N= 5) nmol/mg protein, respectively, significantly different at p< 0.05.

Consistent with our findings of this region being important for CYP4A11/Cyp4a12a-p62 interactions, Cyp4a content as both soluble as well aggregate species was enhanced in 2-month-old p62Mut mouse livers, relative to that in corresponding age-matched WT mouse livers (**Fig. 12B**). Furthermore, this difference in Cyp4a content was held in both freshly isolated and cultured hepatocytes from 2- or 4-month-old p62Mut mice (**Fig. S7**). That this “soluble” Cyp4a stabilization in p62Mut mouse livers is functionally relevant, was documented by liver microsomal ω-hydroxylation of the Cyp4a functional probe arachidonic acid to its physiologically relevant, proinflammatory metabolites 19- and 20-HETEs (**Fig. 12C**), which were significantly (p< 0.0001) increased over the corresponding WT-values (**Fig. 12C**). These findings thus underscore the physiological relevance of p62 in hepatic CYP4A ALD in intact mice.

## DISCUSSION

Our findings described above reveal that in common with other mammalian P450 proteins, CYP4A-family proteins turn over via both ERAD/UPD and/or ERLAD/ALD pathways (14, 15 *and references therein*). However, human liver CYP4A11 and mouse liver Cyp4a12a exhibit an apparent preference for the ERLAD/ALD pathway, as documented by their relative stabilization upon treatment with diagnostic ALD pathway inhibitor probes as well as upon genetic KO of key autophagic initiation proteins ATG5 and BECN1 in mice and/or cell-lines. Such cellular disruption of ERLAD/ALD pathways, led to the stabilization of these P450 proteins in their “soluble” form, readily detected upon extraction with common detergent-containing cell lysis-buffers routinely used for Western IB analyses. However, more surprisingly upon such cellular disruption of ERLAD/ALD pathways, these CYP4A/Cyp4a proteins also accumulated as cellular aggregates that were intractable to common cell lysis buffers routinely used in the extraction of P450s for Western IB analyses. These CYP4A/Cyp4a-containing cellular aggregates required much stronger protein denaturation/solubilization such as boiling in 8 M urea/2 M thiourea-containing TISO buffers for their detection by IB analyses. It is notable however, that to a lesser extent such native CYP4A protein accumulation as aggregate species is also detected in intact (physiologically undisrupted) native, untreated mouse livers (**Fig. 2C**, **Fig. 12**). This finding thus suggests that the total hepatic P450 content of these CYP4A proteins would be normally underestimated and unaccounted for upon routine Western IB analyses of liver microsomes/lysates. The nature of these CYP4A protein aggregates is unclear but may consist of both ubiquitinated proteins as well as protein aggregates destined as cargos for autophagic engulfment and lysosomal clearance. Indeed, it has been proposed (28–30) that in contrast to soluble protein species that are usually destined to ERAD/UPD, such cellular protein aggregates are partitioned to ERLAD/ALD pathways (28–30).

The relative preference of CYP4A11 and Cyp4a12a for turnover via ERLAD/ALD pathway, coupled with our findings of the relative stabilization of these P450 proteins in autophagic receptor (p62/NBR1/OPTN/NDP52/TAX1BP1)-deficient “Penta KO” HeLa cells as well as p62-KO, and p62/NBR-1 double KO HepG2 cells versus in the corresponding WT- or NBR1-KO HepG2 cells, underscored an intimate role for the highly abundant cytoplasmic p62/SQSTM-1 multifunctional and multiprotein scaffold as the relevant autophagic receptor in this process. The identification of the cytoplasmic autophagic receptor p62 as a requirement in CYP4A ALD, in addition to all the other autophagic participants, clearly distinguishes this process from the typical mammalian ER-phagy involving largely ER-resident receptors such as FAM134B, RTN3L, SEC62, CCPG1, ATL3 and TEX264 (6). In this context, it is noteworthy that not only p62 has been shown to be associated with the ER, but also upon withdrawal of the ER-membrane proliferator 1,4-bis[2-(3,5- dichloropyridyloxy)] benzene (TCPOBOP), excess ER as well as TCPOBOP-inducible Cyp2b10 accumulate to a much greater extent in p62-KO mice than in the corresponding WT mice, documenting a role for p62 in ER-homeostasis (76).

This finding also implied the involvement of intimate interactions between the cytoplasmic p62 and the integral ER-anchored CYPs 4A-proteins, which would need to be extracted from the ER-membrane prior to their ALD. Given the presence of ubiquitinated CYP4A11 and Cyp4a12a species in both soluble as well as aggregate states, we considered that plausibly, the p62-UBA- subdomain could be involved in the p62-recruitment of CYP4A11/Cyp4a12a as target substrates. However, both gross and fine structural deletion analyses excluded this plausible p62 UBA-subdomain but revealed that the p62-subdomain relevant to its CYP4A11/Cyp4a12a-interactions was instead confined to its N-terminal 170-233 residues that normally comprise the p62 “LB/TBS”- subdomains (**Fig. 9**). Surprisingly however, neither the deletion of residues 180-189, nor the Ala-mutation of the positively charged R_183_/R_186_/K_187_/K_189_ patch within this specific p62-subdomain affected its CYP4A11/Cyp4a12a-interactions (**Fig. 9B**). This patch was recently identified as a novel IκBα-interaction site (60), as well as an ALS-mutant SOD1-interaction site (56).

Even more surprisingly, p62-oligomerization via its N-terminal PB-1 subdomain, generally required for autophagic cargo presentation for ERLAD/ALD (43–49, 77), apparently was not relevant for its CYP4A11/Cyp4a12a interaction, as the deletion of this PB-1 subdomain failed to affect p62-CYP4A interactions. Interestingly, the p62-subdomain demarcated by 170-256 residues has also been documented to interact with Trim5α, a protein that acts both as an autophagic receptor and an autophagic platform for the assembly of the active autophagy initiation ULK1-Beclin 1 complexes (78). This p62-Trim5α-interaction was likewise found to be independent of the PB-1 subdomain. Intriguingly, an N-terminally spliced p62-variant (Δ1-84 residues), the p62-H2 isoform has been recently identified in the human liver, as an equally if not more abundant isoform than the parent p62-H1 species (58). This p62 isoform, albeit lacking the PB-1 subdomain does tend to form large size aggregates that colocalize with keratin 8 and exhibit a cross-β-sheet conformation characteristic of hybrid inclusions/Mallory-Denk bodies (MDBs) hallmarks of many human liver aggregation diseases (79–83). Our results suggest that this p62-H2 isoform, containing the CYP4A-relevant LB/TB-subdomains, would be also capable of interacting with CYP4A11/Cyp4a12a.

Although, we have identified the p62 hotspots for its CYP4A11/Cyp4a12a-interaction, the corresponding p62-interacting CYP4A regions remain to be identified. Because to our knowledge, P450s are not quite as amenable to similar structural deletion analyses, we have employed *in-cell* chemical XLMS analyses in an attempt to identify these interacting sites. Studies employing the cell-permeable chemical crosslinker DSS in HEK293T cells co-expressing p62 and CYP4A11, indicated that the two proteins clearly interact *intracellularly* (**Figs. 11 & S6**). However, these p62/CYP4A11 complexes albeit clearly detectable upon SDS-PAGE/IB analyses, being highly hydrophobic require very high urea (8M) concentrations for solubilization, which are incompatible with efficient enzymatic trypsin/lysyl endopeptidase C-mediated peptide digestion and/or further enrichment of the crosslinked peptides required for efficient XLMS analyses. To date, these issues have defied our exhaustive attempts to identify the precise p62-interacting CYP4A-sites upon extraction of the complexes in a sufficiently soluble form that can be subjected to peptide mapping and subsequent successful XLMS analyses of the crosslinked peptides.

Nevertheless, inspection of CYP4A11 and Cyp4a12a primary sequences as well as homology models based on the structure of rabbit CYP4B1 (62, 63) reveal the presence of a few surface-exposed LIR-motifs that would be accessible to LC3-interactions required for their autophagosomal targeting (**Fig. 6**). Our co-transfection/co-IP studies with the autophagic marker GFP-LC3 indeed verified its robust interactions with CYP4A11 as well as weaker ones with Cyp4a12a (**Fig. 6**). This suggests that the CYP4A LIR motifs could serve to recruit these P450s directly to the autophagosome, particularly under protein overexpression conditions when their relative pools would dominate over those of other intracellular LIR-rich proteins. However, this may not be the case under physiological conditions when these P450s may need to compete simultaneously with a myriad other LIR-rich cellular proteins for the autophagosomal LC3II-targeting. Notably, while LIR motifs are essential for autophagic targeting, they may not be sufficient. This may explain why specific p62 interactions are normally required for CYP4A-ALD. p62 may insure CYP4A ALD first by marking these P450s as specific autophagic-cargo pool i.e. inclusions/aggregates, and then by engaging LC3 through its own intrinsic LIR-motif, to efficiently target them to the autophagosome for eventual autolysosomal disposal. Not surprisingly, disruption of p62-Cyp4a interactions as observed in the p62Mut mouse livers, results in adefective Cyp4a ERLAD/ALD process, and consequently in the higher level of both soluble and Cyp4a-aggregates (**Figs.12 and S7**).

Collectively, our studies indicate that human and mouse liver CYPs 4A are not only preferential targets of ERLAD/ALD process, but also are recruited to this process by the autophagic receptor p62, through an unprecedented mode of interaction. Disruption of this degradation process at any stage, can result in the stabilization of the CYP4A proteins as functionally active soluble species as well as cellular aggregates, both outcomes of significant physiological and pathophysiological relevance. On one hand, enhanced functional CYP4A levels would lead to the enhanced ω-hydroxylation of its physiological substrate arachidonic acid (35–42) with consequent over generation of 20-HETE, a physiologically active metabolite implicated not only in the regulation of blood pressure, but also in various other pathological processes (inflammation, diabetes, cardiovascular diseases, and cancer) (39, 41, 84–87). On the other hand, the enhanced accumulation of insoluble hepatic CYP4A aggregates would contribute to the pathological deposition of p62-associated Mallory-Denk bodies (MDBs) and other hybrid inclusions, with consequent disruption of the normal hepatic physiological milieu and function (79–83). We have previously reported that the p62Mut mice exhibit an age-dependent liver inflammatory process that we attributed to the observed enhanced NF-κB-transcriptional activation, resulting from impaired p62-mediated protein stabilization of the NF-κB-inhibitor IκBα (60). We now document that such a proinflammatory scenario could undoubtedly also be further exacerbated by the elevated hepatic 20-HETE levels as well as the pathological hepatic MDB-deposition.

## Acknowledgments

We thank Mr. Chris Her for liver cell isolation at the UCSF Liver Center Core on Cell & Tissue Biology, supported by NIDDK Center Grant DK26743. We also thank Dr. N Mizushima (Univ. of Tokyo) and Dr. J. Debnath (UCSF) for providing the ATG5 fl/fl mice, and Dr. Richard J. Youle (NINDS/NIH) for the “Penta-KO” HeLa cells.

## ABBREVIATIONS USED

ALD: Autophagic lysosomal degradation
ATG: Autophagy-related genes
ATG5: autophagy-related 5
BTZ: bortezomib
Co-IP: co-immunoprecipitation
CHX: cycloheximide
P450s; CYPs: cytochromes P450
DMEM: Dulbecco’s Modified Eagle high glucose medium
DSS: disuccinimidyl suberate
ER: endoplasmic reticulum
ERAD: ER-associated degradation
ERLAD: ER-Lysosomal-associated degradation
flp: floxed
HA: hemagglutinin
HMM: high molecular mass
IB: immunoblotting
IB: Immunoblotting
Keap1: Kelch-like ECH associated protein 1
KIR: Keap1-interacting region
KO: knockout
Lys-C: lysylendopeptidase C
LIR: LC3-interacting region
LB: LIM-domain interacting region
3-MA: 3-methyladenine
MEM: minimal Eagle’s medium
LC3: microtubule-associated protein light chain 3
p62 flp/flp: p62-floxed mouse
p62-Myc: Myc-tagged p62
p62mut: p62 genetic mutant mouse
PB: PEST1 and PEST2 motifs phenobarbital
PB1: Phox and Bem1p-domain
SQSTM-1/p62: Sequestosome 1
SOD1: Cu-Zn superoxide dismutase
TBS-subdomain: Traf6-binding
Ub: ubiquitin
UPD: Ub-dependent proteasomal degradation
WT: wild-type
Wort: Wortmannin
UBA: Ub-association
XLMS: chemical crosslinking mass spectrometry
ZZ: Zn-finger binding motifs

## SUPPORTING INFORMATION

**Table S1:**
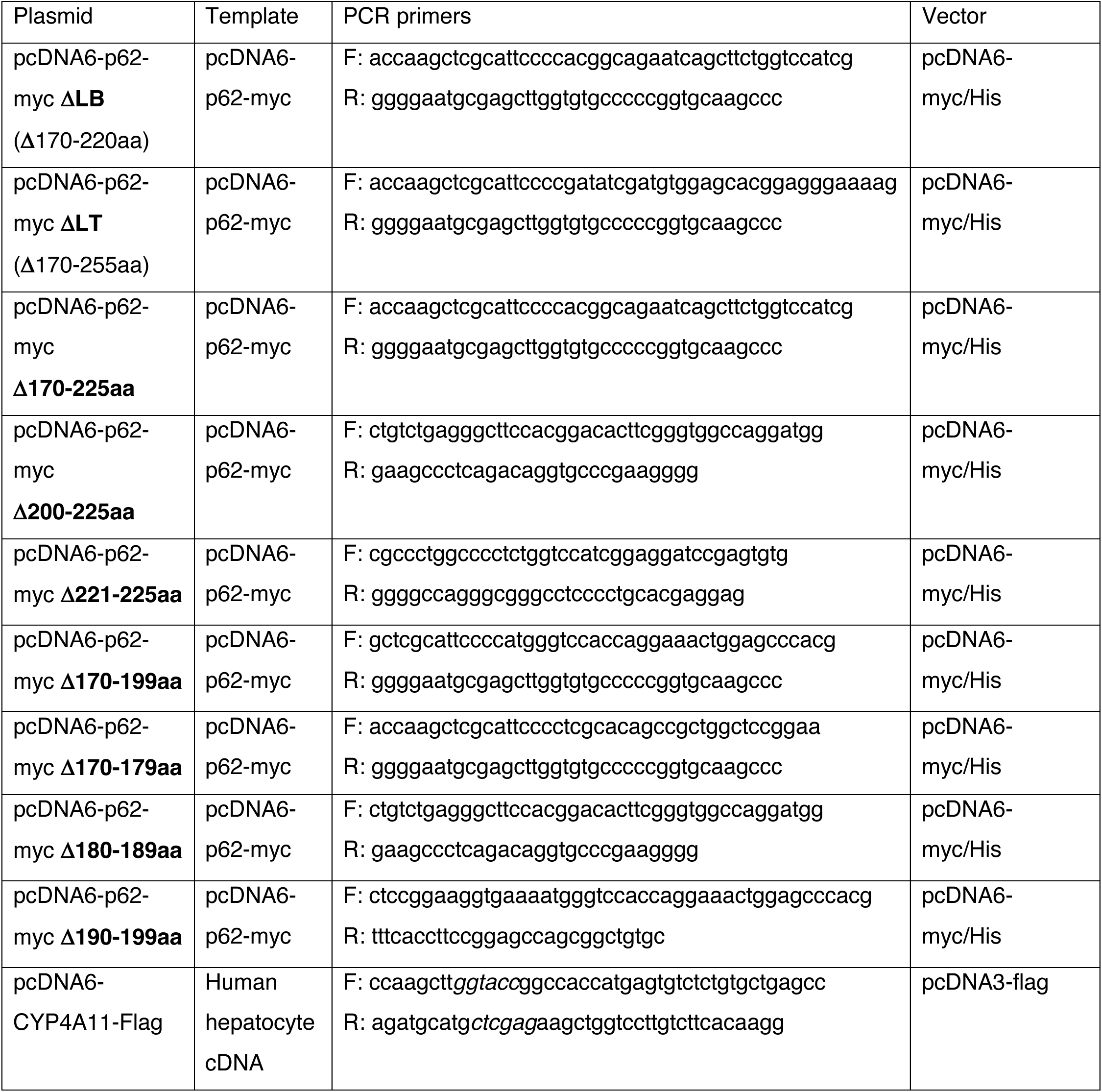
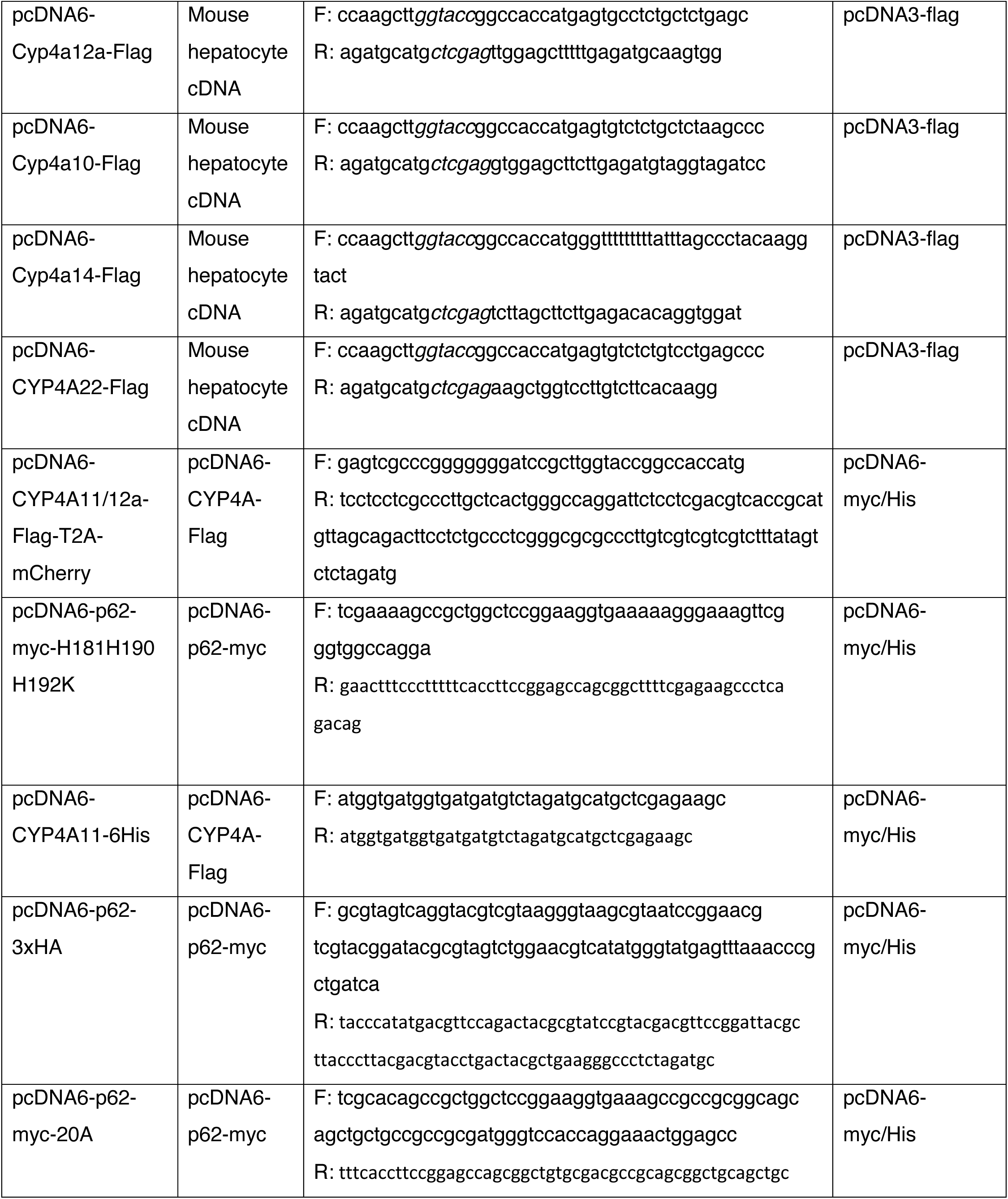
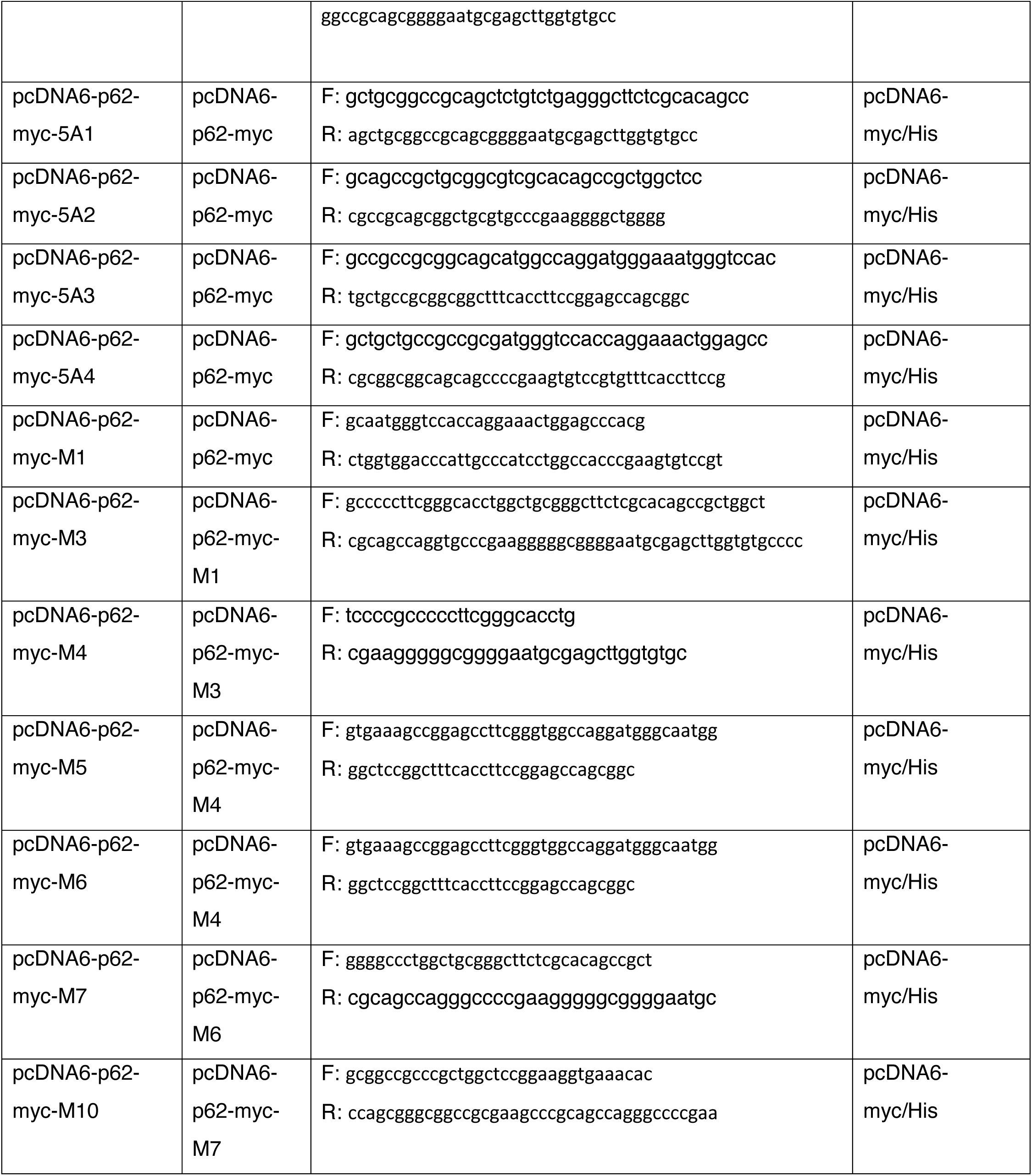
Newly constructed primers, templates, vectors.

**Table S2.**
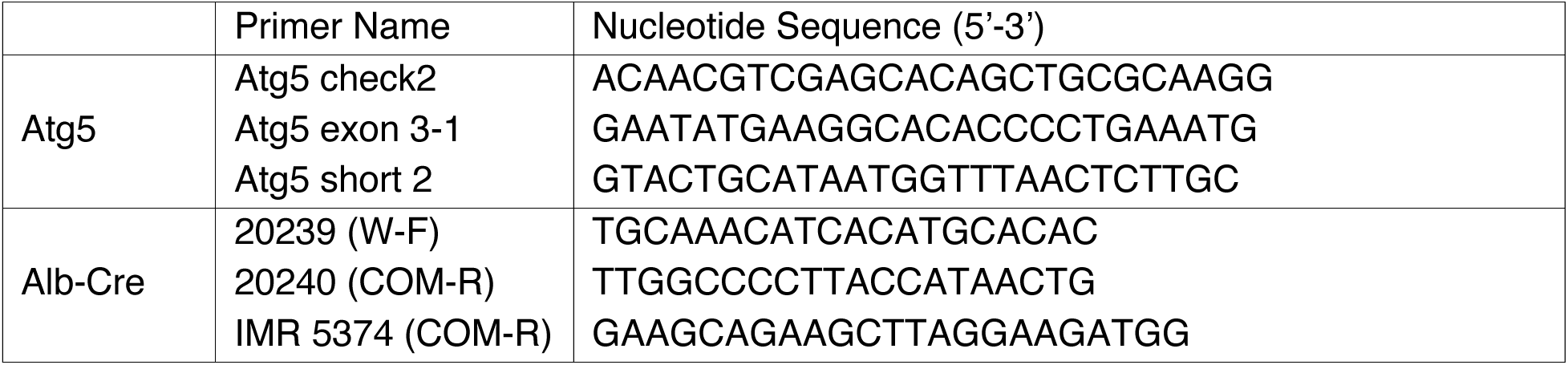
Primer sequences used for Atg5-mouse tail-clip genotyping:

**Table S3.**
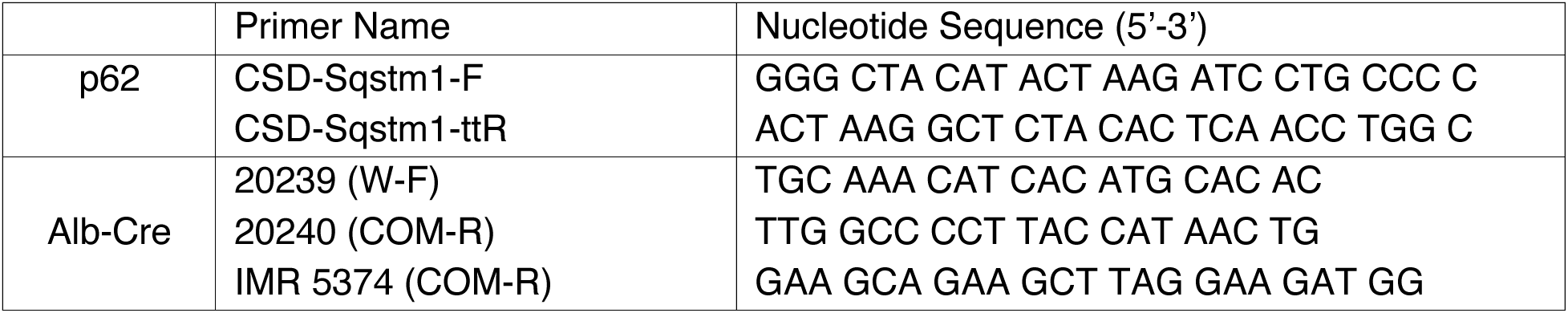
Primer sequences used for p62-mouse tail-clip genotyping:

**Table S4:**
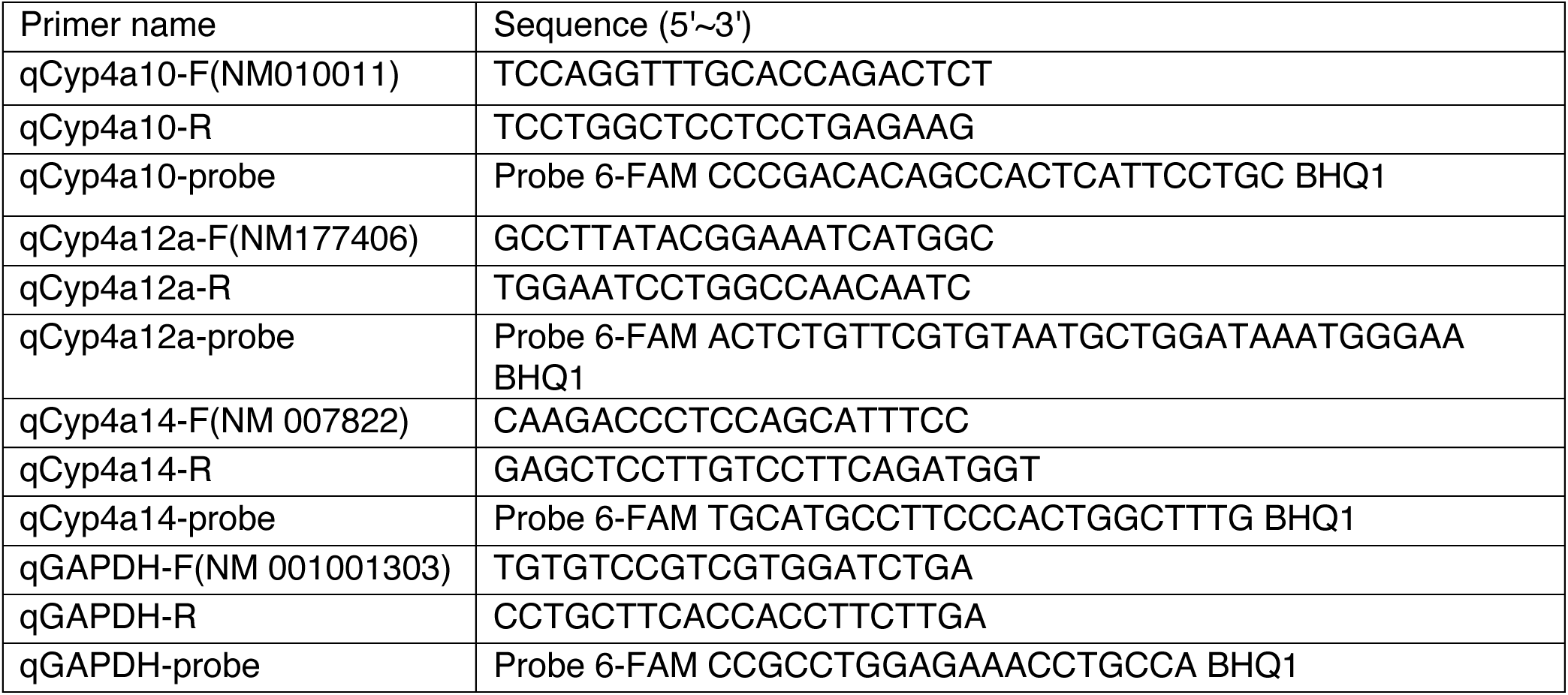
qPCR primers used:

**Fig. S1.**
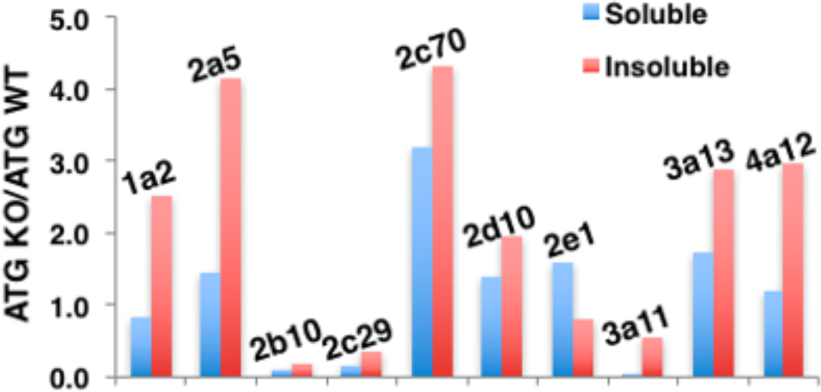
Relative P450 content of Triton-soluble & insoluble fractions upon LC-MS/MS analyses of PB-treated ATG5 KO & WT mouse hepatocytes: Our proteomic analyses of 1% Triton X-100-solubilized and cleared lysates from phenobarbital (**PB**)-pretreated ATG5^-/-^ (knockout; **KO**) and ATG5^+/+^ (wild type; **WT**) 6-week-old mouse hepatocytes, revealed high mass spectral P450 peptide count ratios (KO/WT= >1) revealing the stabilization of certain mouse hepatic P450s upon ALD-disruption. Similar analyses of corresponding insoluble 14,000g pellets of these Triton-solubilized lysates upon further solubilization with TISO buffer revealed not only the parent P450 species, but also their high molecular mass (**HMM**) ubiquitinated/oligomerized aggregates with even higher KO/WT mass spectral peptide count ratios.

**Fig. S2.**
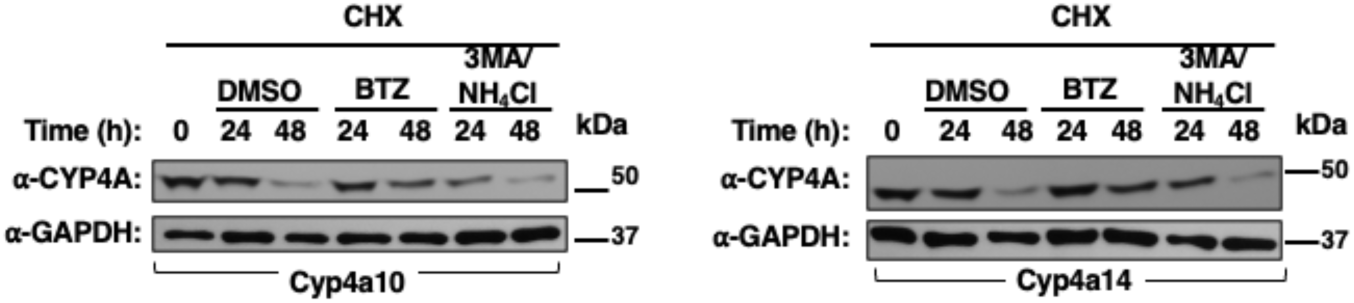
Cyp4a10 and Cyp4a14 are preferentially degraded through UPD rather than ALD in HepG2 cells: HepG2 cells were seeded in 12-well plates overnight, and then each well was transfected with either Cyp4a10-Flag (**A**) or Cyp4a14-Flag (**B**). Forty-eight h after transfection, cells were treated with CHX (50 μg/mL) for 10 min, and then the UPD-inhibitor BTZ, and ALD-inhibitors 3MA/NH_4_Cl were added to specific wells, in parallel. Cells were harvested at indicated times (0, 24 and 48 h), and cell lysates (10 μg) subjected to IB analyses with CYP4A11 antibody, with GAPDH employed as the loading control. Representative immunoblots are shown.

**Fig. S3.**
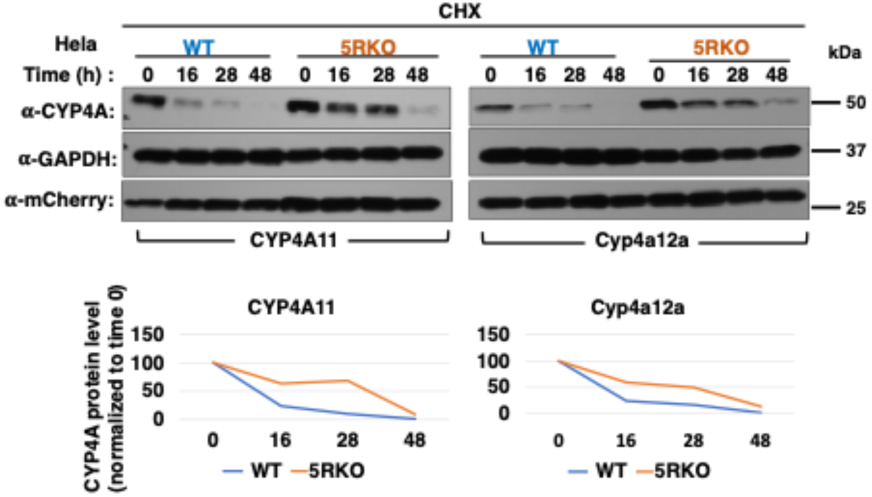
CYP4A proteins are stabilized upon 5RKO in Hela cells: WT Hela cells, and Hela cells with KO of the five autophagy receptor genes (5RKO/pentaKO: TAX1BP1, NDP52, NBR1, p62 and OPTN) cells were seeded in 12-well plates overnight, and then each well was transduced with ssfv-lenti-CYP4A11-Flag-T2A-mCherry or ssfv-lenti-Cyp4a12a-Flag-T2A-mCherry. Forty-eight h after transduction, cells were treated with cycloheximide (CHX; 50 μg/mL). Cells were harvested at indicated times (0, 16, 28, and 48 h). Cell lysates (10 μg) were subjected to IB analyses with CYP4A11 antibody, with GAPDH as the loading control and mCherry as a transduction control. CYP4A amounts relative to 0 h control were quantified from 3 experimental replicates and plotted. The half-life (t_1/2_) of CYP4A in WT and 5RKO Hela cells was calculated based on a single exponential fit of the data with Graphpad Prism Version 6.07. The t_1/2_-values were found to be as follows: CYP4A11 WT (10.2h) and 5RKO (33h); Cyp4a12a WT (11h) and 5RKO (25.4h).

**Fig. S4.**
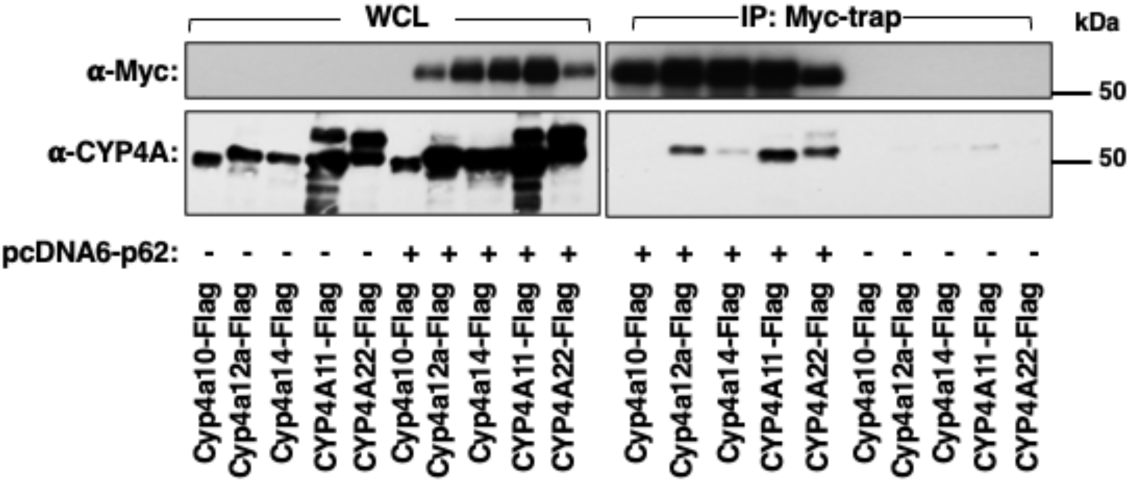
Interactions of Flag-tagged human CYPs 4A and mouse Cyps 4a with p62-Myc: HEK293T cells were seeded in 60 mm dishes overnight, and then each dish was co-transfected with pCDNA3.1-p62-myc (1 μg) and Flag-tagged CYPs 4A or Cyps4a (4 μg) for 48 h. Cells were treated with Torin(1μM) together with MG262 (1nM)/NH_4_Cl (30 mM) 24h prior to the cell harvest. The cells were harvested with cell lysis buffer containing Triton X-100, protease and phosphatase inhibitor cocktail, and 0.1% SDS; Whole cell lysates (500 μg) were used for co-IP with Myc-trap (50 μl), followed by IB analyses with CYP4A11- and Myc-antibodies. IP, coimmunoprecipitation; IB, immunoblotting; WCL, whole cell lysates.

**Fig. S5.**
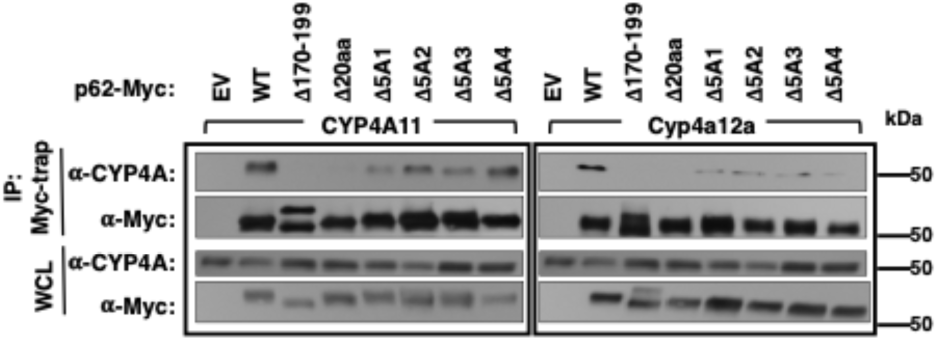
Identification of p62-CYPA11/Cyp4a12a-interaction domains through finer structural-deletion analyses of p62/SQSTM-1 170-179 and 190-199 subdomains: p62 subdomain deleted mutants were constructed as indicated (EXPERIMENTAL PROCEDURES). HEK293T cells were seeded in 60 mm dishes overnight, and then each dish was co-transfected with pcDNA3.1-CYP4A11 (4 μg) or Cyp4a12a (4 μg) and either WT or indicated mutant pcDNA6-p62-Myc vector (2 μg) for 48 h, and treated exactly as indicated in Fig.7. Whole cell lysates (WCL) (500 μg) were used for co-IP with Myc-trap (50 μl), followed by IB analyses with CYP4A11 and Myc-antibodies. IP, coimmunoprecipitation; IB, immunoblotting; EV, empty vector; WT, p62 WT vector; Δ170-199, p62 with deletion of 170-199 subdomain; Δ20aa, p62 with deletion of 170-179 and 190-199 subdomains; Δ5A1, Δ170-174, deletion of 170-174 p62-subdomain; Δ5A2, Δ175-179, deletion of 175-179 p62-subdomain; Δ5A3, Δ190-194, deletion of 190-194 p62-subdomain; Δ5A4, Δ195-199, deletion of 195-199 p62-subdomain.

**Fig. S6.**
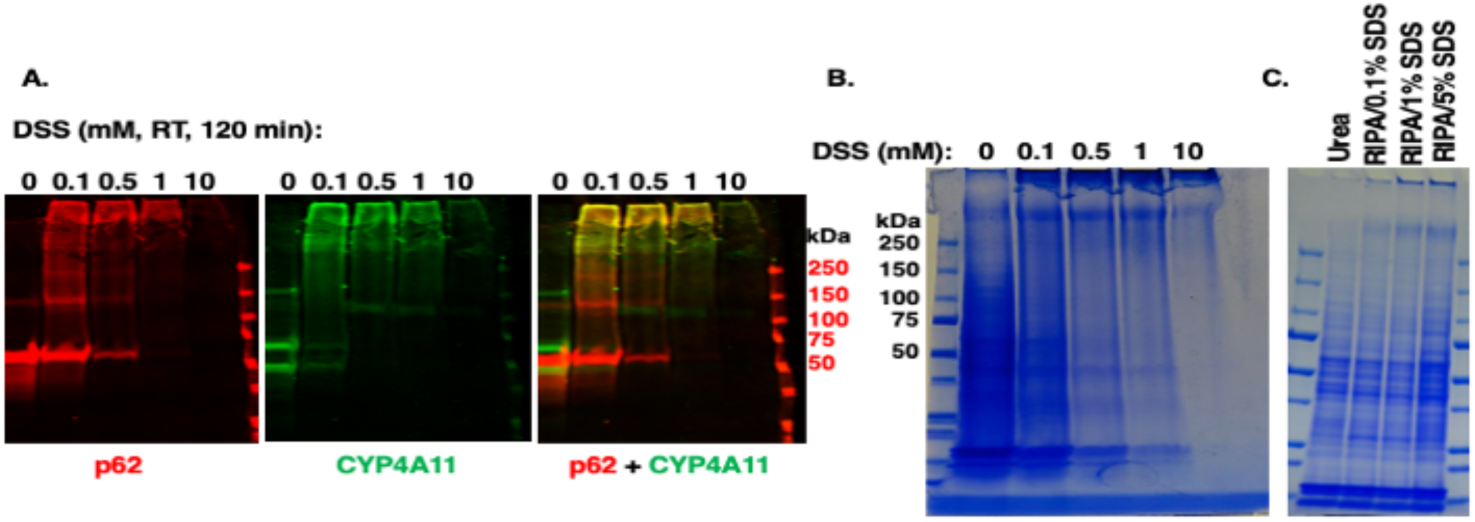
Preliminary optimization of in cell DSS-elicited crosslinking conditions and urea/SDS-concentrations required for solubilization of the crosslinked CYP4A11 and p62-HA: **A.** HEK293T cells overexpressing CYP4A11[His]_6_ and p62[HA]_3_ were treated with DSS *in cell* at the indicated concentrations, lysed in RIPA buffer, and then immunoblotted with rabbit anti-HA and mouse anti-His antibodies Licor staining. **B**. The samples from A were used for the SDS-PAGE/Coomassie Bíilliant Blue staining. **C**. HEK293T cells overexpressing CYP4A11[His]_6_ and p62[HA]_3_ were treated with DSS (0.1 mM) at room temperature for 2 h in the dark, and the cells were lysed with 8 M urea, and RIPA buffer containing SDS at the indicated concentrations, followed by SDSPAGE/Coomassie Bíilliant Blue staining.

**Fig. S7.**
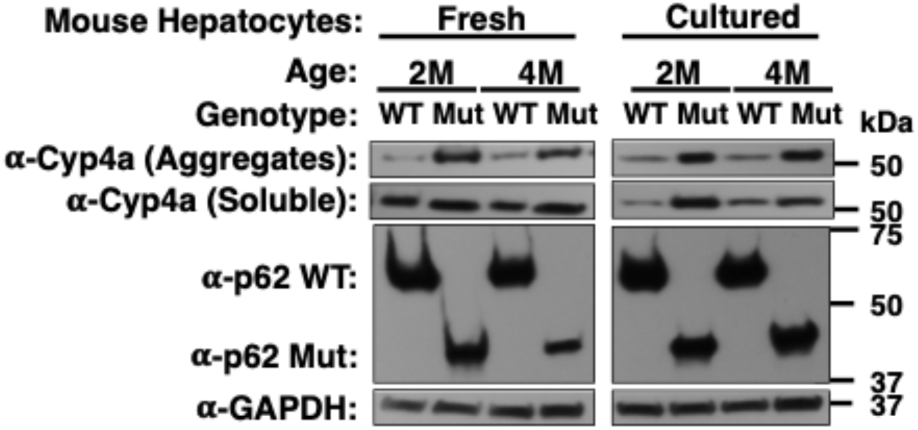
Relative Cyp4a stabilization in p62mut vs WT mouse cultured hepatocytes: p62 Western IB analyses of hepatocytes isolated from 2-month- or 4-month-old p62 WT and p62Mut mice. Hepatocytes were homogenized in cell lysis buffer and sedimented at 14,000g to obtain the insoluble CYP4A aggregates. The pellet containing CYP4A aggregates was solubilized in TISO buffer as described (Experimental Procedures). Cell lysates or solubilized pellet (10 μg) were subjected to IB analyses with CYP4A11 antibody, use GAPDH as the loading control.

